# LSD1 inhibition corrects dysregulated MHC-I and dendritic cells activation through IFNγ-CXCL9-CXCR3 axis to promote antitumor immunity in HNSCC

**DOI:** 10.1101/2025.03.17.643710

**Authors:** Amit Kumar Chakraborty, Lina Kroehling, Rajnikant Dilip Raut, Chumki Choudhury, Maria Kukuruzinska, J Silvio Gutkind, Xaralabos Varelas, Bikash Sahay, Stefano Monti, Manish V. Bais

## Abstract

Poor infiltration of CD8+ T cells and dysregulated MHC-I confer resistance to anticancer clinical therapies. This study aimed to elucidate the mechanisms of lysine-specific demethylase 1 (LSD1, encoded by KDM1A gene) in antitumor immunity in Head and Neck Squamous cell carcinoma (HNSCC). LSD1 inhibition in syngeneic and chronic tobacco carcinogen-induced HNSCC mice recruited activated dendritic cells (DCs), CD4+ and CD8+ T cells, enriched interferon-gamma (IFNγ) in T cells, CXCL9 in DCs, and CXCR3 in T cells, as evaluated using flow cytometry and single-cell RNA-seq analysis. Humanized HNSCC mice and TCGA data validated the inverse correlation of KDM1A with DC markers, CD8+ T cells, and their activating chemokines. *Kdm1a* knockout in mouse HNSCC and LSD1 inhibitor treatment to co-culture of human HNSCC cells with human peripheral blood mononuclear cells (PBMCs) resulted in MHC-I upregulation in cancer cells for efficient antigen presentation in tumors. Overall, LSD1 inhibition in tumor cells upregulates MHC class I and induces DCs to produce CXCL9, which in turn activates CD8+ T cells through the CXCL9-CXCR3 axis to produce IFNγ. Finally, we identified a novel mechanism by which LSD1 inhibition promotes the activation of H3K4me2 and its direct interaction with MHC-I to induce antitumor immunity. This may have implications in poorly immunogenic and immunotherapy-resistant cancers.

**Statement of Significance:** LSD1-mediated unique mechanisms have impact on epigenetic therapy, MHC-I resistant HNSCC therapies, and poor CD8+ and dendritic cell infilterated tumors.

## Introduction

Head and neck cancer (HNC) is associated with significant mortality, with a survival rate of less than 50%. We showed that inhibition of the epigenetic regulator lysine-specific demethylase 1 (LSD1) reduces tumor progression (1) and promotes infiltration of CD8+ T cells (2,3). However, the role of LSD1 in promoting <u>h</u>ead and neck <u>s</u>quamous <u>c</u>ell <u>c</u>arcinoma (HNSCC)-specific mechanisms of antitumor immunity is not well understood and could offer new directions for treatment. A recent study has shown that an intratumoral immune triad comprising CD4+, CD8+, and dendritic cells (DCs) is required for efficient solid tumor elimination (4).

DCs (CD11c+ MHC class II+) have been grouped into categories based on their role and specific marker expression, such as conventional DC type I or cDC1 (XCR1+), conventional DC type 2 or cDC2 (CD11b+), and migratory DC or mDC (CD103+ or CCR7+). cDC1 can efficiently present antigens to CD8+ T cells, initiating robust cytotoxic T lymphocyte (CTL) responses critical for tumor elimination (5–8). cDC2 relies on CD4-mediated help signals in immunotherapy (9,10). mDCs transport intact antigens to the lymph nodes and prime tumor-specific CD8+ T cells (11). These migratory DCs are capable of inducing cytotoxic T lymphocytes (CTL)(12).

IFNγ produced by CD8+ T cells and other immune cells can induce the expression of CXCL9 on DCs, which binds to CXCR3 in T cells and recruits more CD8+ T cells to the tumor site. Alternatively, sensing of T cell-secreted IFNγ by bystander tumor cells may potentially boost MHC-I expression by inducing expression CXCL9 and directly inducing tumor cell death (13). The CXCL9/CXCR3 axis regulates immune cell migration, differentiation, activation, and Th1 cell polarization (14). IFNγ shows a vast and homogeneous distribution and reprograms tumor microenvironment (TME) (13). Cancer cells express neoantigens on the cell surface via MHC I and II molecules (15). MHC-I plays a vital role in priming cytotoxic T cells, leading to antitumor responses (16). MHC class II on cDC1 cells is required for CD4+ T cells to facilitate efficient CD8+ T cell priming during antitumor immune control in mice.

Humanized cancer models are suitable for supporting non-human leukocyte antigen (HLA)-matched tumor growth. The presence of human immune cells does not significantly affect tumor growth rates (17). These mice have been extensively used in PD-1 and PD-L1 immunotherapy-related studies (18,19).

The aim of this study was to evaluate the role of LSD1 inhibition in promoting CD8+ T cell infiltration during HNSCC growth. We hypothesized that LSD1 inhibition promotes CD8+ T cells with the support of antigen presentation by DCs and cytokines in patients with HNSCC. To account for variability, we evaluated the mechanism of LSD1 in a syngeneic model, chronic tobacco carcinogen-induced, novel humanized mouse model, *Kdm1a* genetic knockout mouse models, ex vivo co-cultures, single-cell RNA sequencing, and validation by immunophenotyping. Finally, we evaluated how LSD1 inhibition promotes antitumor immunity by activating the novel IFNγ-CXCL9-CXCR3 axis and MHC-I-mediated mechanisms.

## Material and Methods

### Syngeneic 4MOSC1 model

4MOSC1 cells were grown in keratinocyte serum-free media and injected into the tongues of mice (2.5 × 10^5^ cells). Two days post-injection, mice were treated daily with the local application of vehicle, SP2509 high-dose (SH; 40 mg/kg), or vehicle in the respective groups. After 14 days of treatment, the mice were sacrificed and tongue tumor lesions were dissected and used for flow cytometry, qPCR, and RNAseq analysis.

### 4-Nitroquinoline 1-oxide (4NQO1) induced mouse model

All experiments were performed with prior approval from the Institutional Animal Care and Use Committee (BUMC IACUC) at Boston University using male and female mice. C57BL/6J mice were fed 100 μg/mL 4NQO (in propylene glycol) in drinking water for 16 weeks, followed by regular drinking water for the remainder of the study period. Exposure of the tongue epithelia to 4NQO results in early and advanced stages of the disease, including hyperplasia (weeks 0–8), papilloma/dysplasia (weeks 9–18), and HNSCC (weeks 18–25) (20). This model captured pathological changes that were similar to those observed in human HNSCC (21). The mice were treated daily after week 20 with the local application of vehicle, low-dose SP2509 (20 mg/kg), or high dose (40 mg/kg). The mice were sacrificed after week 23 and “tongue tumor lesions,” which could encompass invasive carcinomas as well as dysplasia, were collected, to evaluate the entire lesion with an unbiased approach scRNA-seq, RT-qPCR analyses as well as flow cytometry, histology in the respective groups.

### Humanized HNSCC stem cell mice model

NOD-Prkdc^em26Cd52^Il2rgem^26Cd22^/NjuCrl (NCG) coisogenic, CRISPR genetically engineered immunodeficient mice (Charles River Laboratories), and pre-qualified GVHD human peripheral blood mononuclear cells (PBMCs), which were validated for engraftment, were used to establish humanized mice. NCG mice lack functional T cells, B cells, and NK cells and have reduced levels of DCs and macrophages.

The NCG mice were injected with hPBMC (100,000/mouse) via their tail veins. After three days, authenticated and validated HNSCC stem cells obtained from Cellprogren (cat. #36125-52P, Lot #2111210) (table including marker details provided by Celprogen included in Figure S8) were engrafted in NCG mice (orthotopically engrafted into their tongues (250,000 cells/mouse; n=10/condition)) using our standard protocol (22,23). The mice were treated orally with vehicle or orally administered SP2509 (40 mg/kg, 5 times a week for four weeks) starting from days 3 to 28 post-HNSCC cell engraftment, and the tongues were collected to observe changes. Patient-derived tongue epithelial (PTE) HNSCC primary cells were implanted into the tongue of mice. After seven days, humanized HNSCC mice were treated with SP2509 (40 mg/kg) daily for four weeks, followed by sacrifice and flow cytometry for human immune cells.

### Keratin 14 promoter-specific Kdm1a (LSD1) deficient mice

As shown in our previous study (2), the conditional deletion of Lsd 1 was performed by crossing Lsd1-floxed mice obtained from Stuart Orkin laboratory [Mass General Hospital, Boston (24) with mice expressing Cre recombinase from the Krt14 promoter (25) followed by 4NQO induction and tamoxifen-mediated deletion of *Kdm1a*. *Kdm1a^-/-^* mice were sacrificed after 23 weeks after post-4NQO exposure, and stained tongue tumor sections showed Kdm1a deletion (*Kdm1a^-/-^*). K14 promoter-driven conditional Kdm1a-floxed mice treated with vehicle were designated as *Kdm1a^WT/WT^*, whereas tamoxifen-treated mice with deleted kdm1a were designated as *Kdm1a^-/-^*. Immunofluorescence analysis was performed on n=3/group and 3 sections/mouse, followed by confocal microscopy and image analysis.

### HSC3:PBMC co-culture model

A total of 500,000 HSC3 cells per well were grown in a 6-well plate in DMEM, 10%FBS, and 1% penicillin-streptomycin overnight and treated as follows: 1) vehicle, 2) SP2509 (1 µM), 3) scrambled sgRNA, and 4) Kdm1a sgRNA for another 24 h. The medium was replaced with fresh medium, followed by the addition of 20,000 PBMCs for another 24 h. Finally, the cells were fixed and stained using a flow cytometer. For the CXCL9 and CXCR3 blocking experiments, 1 µg of Human CXCL9 Antibody (R&D Systems # MAB392-SP) and Human CXCR3 Antibody (R&D Systems # MAB160-SP) blocking antibodies per well were added, followed by co-culture with human PBMCs.

### Flow cytometry

A detailed list of all antibodies used is provided in the Supplementary Table. Briefly, CD11c and MHC class II antibodies were used to detect dendritic cells (DCs). XCR1+ CD11c+ MHC II+ cells were used to detect cDC1, CD8a+ CD11c+ MHC II+ cells to detect resident DCs (rDCs), and CD103+ CD11c+ MHC II+ cells to detect migratory DCs (mDCs). CCR7 is another mDC marker(26) used in scRNA-seq studies. The defined T cells included the CD3+TCRβ+ subpopulation of live CD45+ cells for further differentiation into CD4+ (T_H_ cells), CD8+ (cytotoxic T cells), NK1.1+ CD3+ (NKT cells), and NK1.1+ CD3-(NK cells) T cell subtypes. Human CD45 and CD3 were used to identify the leukocytes and lymphocytes, respectively.

### Chromatin Immunoprecipitation (ChIP)

ChIP analysis was performed with ∼4×10^6^ cells using the SimpleChIP® Enzymatic Chromatin IP Kit (Magnetic Beads) according to the manufacturer’s instructions (Cell Signaling Technology, #9003s). Anti-H3K4me2 Recombinant Rabbit Monoclonal Antibody (24H8L19) (Thermo Scientific, #701764) and anti-H3K9me2 Polyclonal Antibody (Thermo Scientific, #39239) for ChIP were used to evaluate their interactions with MHC-I.

### Single-cell RNA-seq

C57BL/6J mice were treated with 4NQO1 for 16 weeks, followed by regular drinking water (no 4NQO1) for the remainder of the experiment. At week 20, these mice were divided into three groups (n=6) and subjected to different treatments as groups-1) vehicle, 2) 4NQO; and 3) 4NQO+SP2509. At the time of sacrifice after week 23, tongue tumor lesions were cut to prepare single-cell suspensions, followed by scRNA-seq analysis of ∼8000 cells/mouse (n=2/condition) using a 10X genomics kit according to the manufacturer’s recommendations. Single-cell suspensions were loaded on a 10X Genomics Chip B and processed using the 10X Genomics 3’v3 kit (10x Genomics, CA, USA). Following the 10X Genomics Chromium controller, the gel beads in emulsion (GEMs) were transferred to strip tubes and subsequently placed into a thermal cycler using the following parameters: 45 min at 53 °C, 5 min at 85 °C, and hold at 4 °C. The samples were then stored overnight at -20 °C. The next day, the samples were thawed at room temperature before the addition of the recovery agent to break the GEMs. Subsequently, cDNA was purified and amplified, and gene expression libraries were generated and purified. The size distribution and molarity of the libraries were assessed using a Bioanalyzer High-Sensitivity DNA Assay (Agilent Technologies, Lexington, MA, USA). The libraries were then pooled at 5nM and sequenced on an Illumina NextSeq 500 instrument at a 1.5-1.9 pM input and 1% PhiX spike in using 150 cycles High Throughput flow cells (Illumina, San Diego, CA, USA) in several sequencing runs resulting in 43,669 - 47,157 mean reads per cell.

### Single-cell RNA-seq data analysis

The data were processed with Cell Ranger v3.1.0 using the mouse reference genome mm10 v2.1.0. SingleCellTK was used to characterize low-quality cells (27). Matrix files were loaded into Seurat v5.0.3 (28) with no initial cutoffs to produce a Seurat object from which low-quality cells were removed. Low-quality cells were defined as those with > 20% reads coming from mitochondrial genes, those with less than 400 genes detected, cells with > 50% contamination as determined by decontX, or cells that were marked as doublets by all scDblFinder metrics (29,30). Filtering reduced the total cells from 76275 to 50319. Data were normalized by library size and log-transformed using normalized data on the RNA slot. Clustering was performed with 40 PCs at a resolution of 1 using 5000 variable features. Cells were scored for the cell cycle, and singleR was used to classify cells at the single-cell level using the Immgen database (31,32). Cell labels were extended to cluster-level labels based on the majority cluster labels and cell marker genes.

The immune cells were a subset and re-clustered using 20 PCs at a resolution of 0.3. Marker genes were used to further define the immune subtypes. Differentially expressed genes were found for each cluster using MAST, with batch as a fixed effect. The dendritic cells were a subset and re-clustered using 20 PCs with a resolution of 0.3. Langerhans cells were removed from downstream analyses because of their distinct transcriptomic profiles. T cells were subsets of the immune compartment by selecting the “T/NK”, “TGD”, and “ILC2” clusters. They were re-clustered using ten PCs at a resolution of 0.4. Differential expression analysis for the Epithelial and T cell compartments (excluding ILC2s due to their distinct transcriptional signature) was performed between treatments in a pairwise manner using MAST, including batch as a fixed effect.

CellChat v2.1.2 was run on the RNA data slot and separately on the 4NQO control and SP2509 treated samples (33). Overexpressed genes were identified using a 1% threshold percentage. Communication probabilities were computed using trim = 0.05, type = truncated mean. Communications were filtered to include those with at least five cells. The two cell chat objects were then merged to compare their interaction occurrence and strength.

### RNA extraction and analysis

TRIzol reagent was used to extract total RNA. RNA-seq and gene set enrichment analyses were performed as described in our previous studies (34) using 400 ng of total RNA for sequencing with Novoseq (35,36). Raw FASTQ sequencing reads were mapped to the M. musculus reference genome (mm10). Differential gene expression analysis was performed using DESeq2 in the R/Bioconductor software.

### Pathological characterization and immunostaining

H&E staining and immunofluorescence were performed on n=4/group and four sections from each mice.

Tongue sections were evaluated for pathology by a board-certified pathologist as established earlier (2).

### The Cancer Genome Atlas (TCGA) analysis

TCGA data were obtained using the TCGABiolinks R/Bioconductor package. Immune infiltration analysis data were acquired from the TIMER 2.0, web portal(37).

### Ingenuity Pathway Analysis

Ingenuity Pathway Analysis was performed using IPA software (Qiagen) (38,39). We have used RNAseq differential gene expression (DGE) data generated with DESeq2 (R/Bioconductor software) to perform the analysis. The genes were included with the criteria using FDR value <0.05 and Z score ≥ ±0.25.

### Statistical analysis

Data analysis for all experiments was performed using the Brown-Forsythe and Welch ANOVA tests.

### Data access statement

All relevant data are within the paper and its Supporting Information files.

### Ethics statement

The study was approved by the BU IACUC committee.

## Results

### The LSD1 inhibitor (SP2509) promoted the accumulation of CD4+, CD8+ T cells, and DCs in a syngeneic HNSCC mouse model

To evaluate whether LSD1 (*Kdm1a*) play a role in tumor immunity, a syngeneic 4MOSC1 mouse model was used (Figure 1A). Treatment with a small-molecule LSD1 inhibitor SP2509 reduced the tumor volume (Figure 1B, Figure S1A) and Kdm1a expression (Figure 1C). Flow cytometry analysis showed that the SP2509 treated group had a significant accumulation of immune cells (Figure 1D, E), especially CD4+ (p<0.01) and CD8+ (p<0.001) T cells, NK cells (p<0.001), and NKT cells (p<0.001), compared with the vehicle treatment group. Analysis of antigen-presenting cells (APC) showed that SP2509 treatment also recruited CD8a+ DC (resident DC) (p<0.01), CD103+ DC (migratory DC) (p<0.001), and XCR1+ DC (Conventional DC1 or cDC1) (p<0.01) (Figure 1F, Figure S1B). The results showed that LSD1 inhibition in the 4MOSC1 syngeneic model (which corresponds to pre-existing HNSCC) by SP2509 promoted the infiltration of CD4+ and CD8+ T cells, NK and NKT cells, and DCs, which could play a role in promoting antitumor immunity. However, the analysis of spleen tissues from SP2509-treated mice did not reveal significant differences in these immune populations (Figure S1C-E).

**Figure 1:**
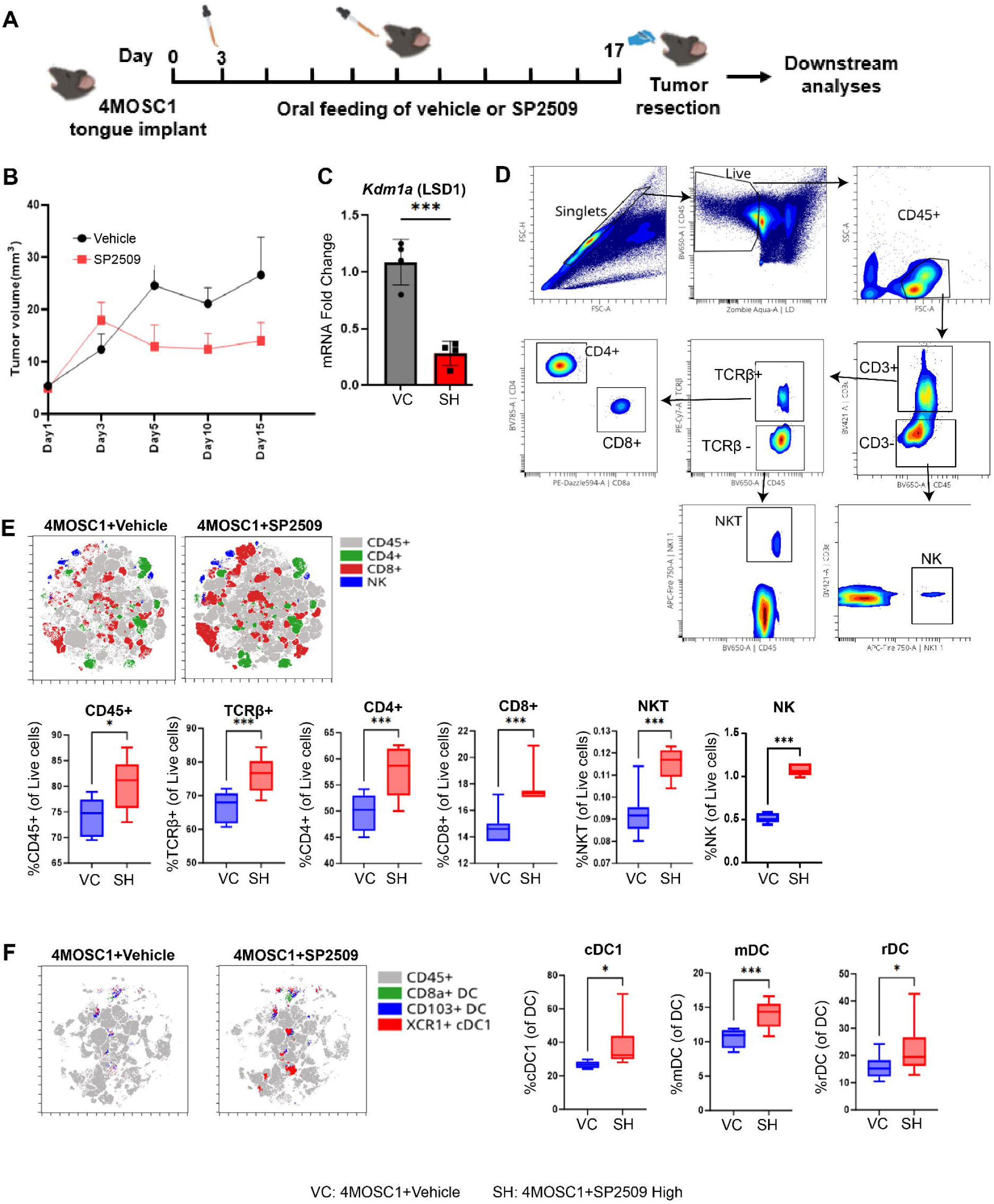
LSD1 inhibition by local SP2509 promoted infiltration of DCs, CD4+, and CD8+ T cells in HNSCC. A) Experimental design showing tobacco carcinogen-induced 4MOSC1 primary cells implantation into the tongue of C57BL/6J mice. Post-3-day implantation, SP2509 in corn oil was applied five times a week starting from day 3-17, followed by multicolor flow cytometry and RNA sequencing. B) Measurements of tumor volume at 3 to 5-day intervals for vehicle or SP2509 treated mice. C) Analysis of Kdm1a (LSD1) expression levels using quantitative real-time polymerase chain reaction (RT-qPCR). Comparing vehicle control (VC) and SP2509high (40mg/kg) (SH) groups using 2^(-ΔΔCT)^ method. D) Gating strategy for detecting immune cell surface markers using multicolor flow cytometry; E) t-SNE and box plots representing higher levels of immune cells in SP2509 treated mice. F) t-SNE and box plots representing the higher levels of specialized dendric cells [cDC1 (XCR1+ CD11c+ MHC class II+), mDC (CD103+ CD11c+ MHC class II+), and rDC (CD8α+ CD11c+ MHC class II+)] in vehicle or SP2509 treated mice. (*p<0.05, **p<0.01, ***p<0.001, ****p<0.0001).

### LSD1 inhibition promoted activation of CD4+, CD8+ T cells, and DCs in chronic 4NQO1 mouse HNSCC

To understand the progressive mechanism and reproducibility of the above study, and to better understand the activation and effector function of T cells and DCs, a chronic tobacco carcinogen 4NQO1-induced HNSCC model was used (Figure 2A). 4NQO1 promotes the transformation of normal tongue epithelium through stage-specific mutations similar to those in human HNSCC. SP2509 treated mice showed a dose-dependent decrease in *kdm1a* expression (Figure 2B), tumor pathologically invasive lesions (Figure 2C), and quantification of invasive pathological lesions (Figure 2D). As shown in Figure 2E, SP2509 treatment resulted in an increase in CD45+, TCRβ+, CD4+, CD8+, and NKT cells compared with the vehicle treatment group (Figure S2A, B). Next, we evaluated activation (CD69+) and effector function (CD44+) markers in conjunction. SP2509 treatment promoted the activation of CD4+, CD8+, NKT, and NK cells, which have effector functions (Figure. S2C). However, SP2509-treated mouse spleens did not show significant differences in these immune populations (Figure S3A-C). Overall, SP2509 promoted the infiltration and activation of effector DCs and T cells in tongue tumors.

**Figure 2:**
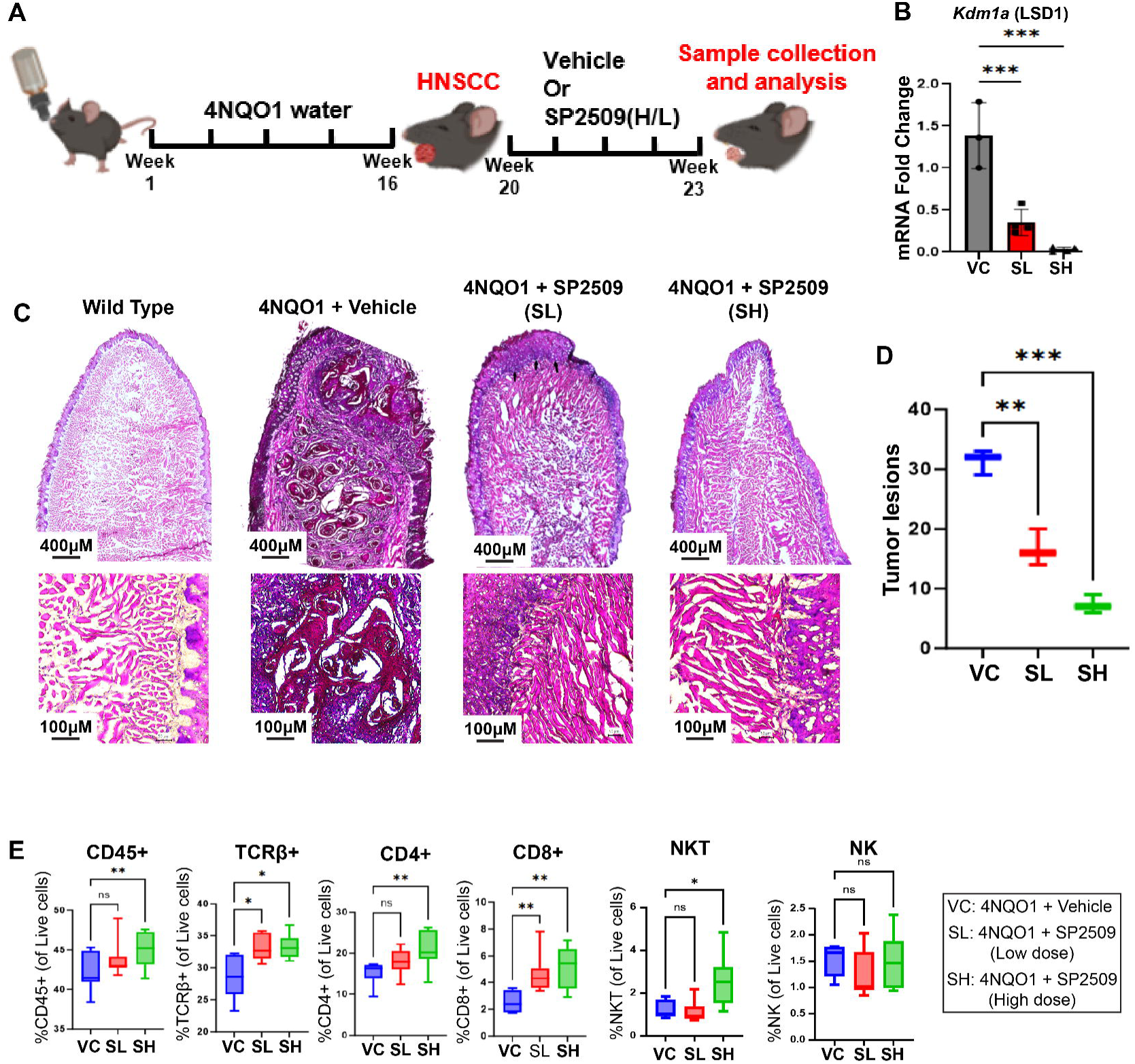
LSD1 inhibition promotes T-cell infiltration in chronic 4NQO-induced progressive HNSCC mouse tongue. A) The experimental design shows that C57BL/6J mice were fed with 4NQO for 16 weeks, followed by regular drinking water (no 4NQO1) for the remainder of the experiment, to develop HNSCC. Vehicle or SP2509 treatment started at week 20 and continued until week 23. B) RT-qPCR showing reduced expression of Kdm1a after treatment with SP2509 low (SL) or SP2509 high (SH) doses. C) H&E staining of mice tongue treated with SL or SH of SP2509. D) Boxplots representing the quantification of tumor lesions per tongue evaluated by H&E-staining. E) Flow cytometry analysis for the characterization of immune cell (T/NK cells) population in mice tongue from SP2509 SL and SH treatment group. (*p<0.05, **p<0.01, ***p<0.001, ****p<0.0001).

### LSD1 inhibition promotes IFNγ, CXCR3 and CXCL9 expression during activation of DCs

To evaluate the mechanism of DC accumulation, we analyzed global transcriptomic changes using bulk RNA-seq analysis of mice harboring tumors treated with or without SP2509. First, we confirmed that SP2509 promoted immune response-related genes, including Ltf, which are related to T cell activation (Figure 3A). GSEA showed that SP2509 treatment promoted positive enrichment of the immune response, innate immune response, defense response, T cell activation, cytokine activity, and cytokine-mediated signaling pathway networks (P<0.05), which could be responsible for CD8+ T cell activation (Figure 3B, C). Ingenuity pathway analysis (IPA) of differentially expressed genes predicted activation of a cytokine network that includes IFNγ, CXCR3, and CXCL9, which has the potential to induce the proliferation of CTLs (Figure 3D). Finally, validation with flow cytometry showed that LSD1 inhibition enriched CXCR3 and IFNγ in activated (CD69+) CD4+ and CD8+ cells, respectively (Figure 3E).

**Figure 3:**
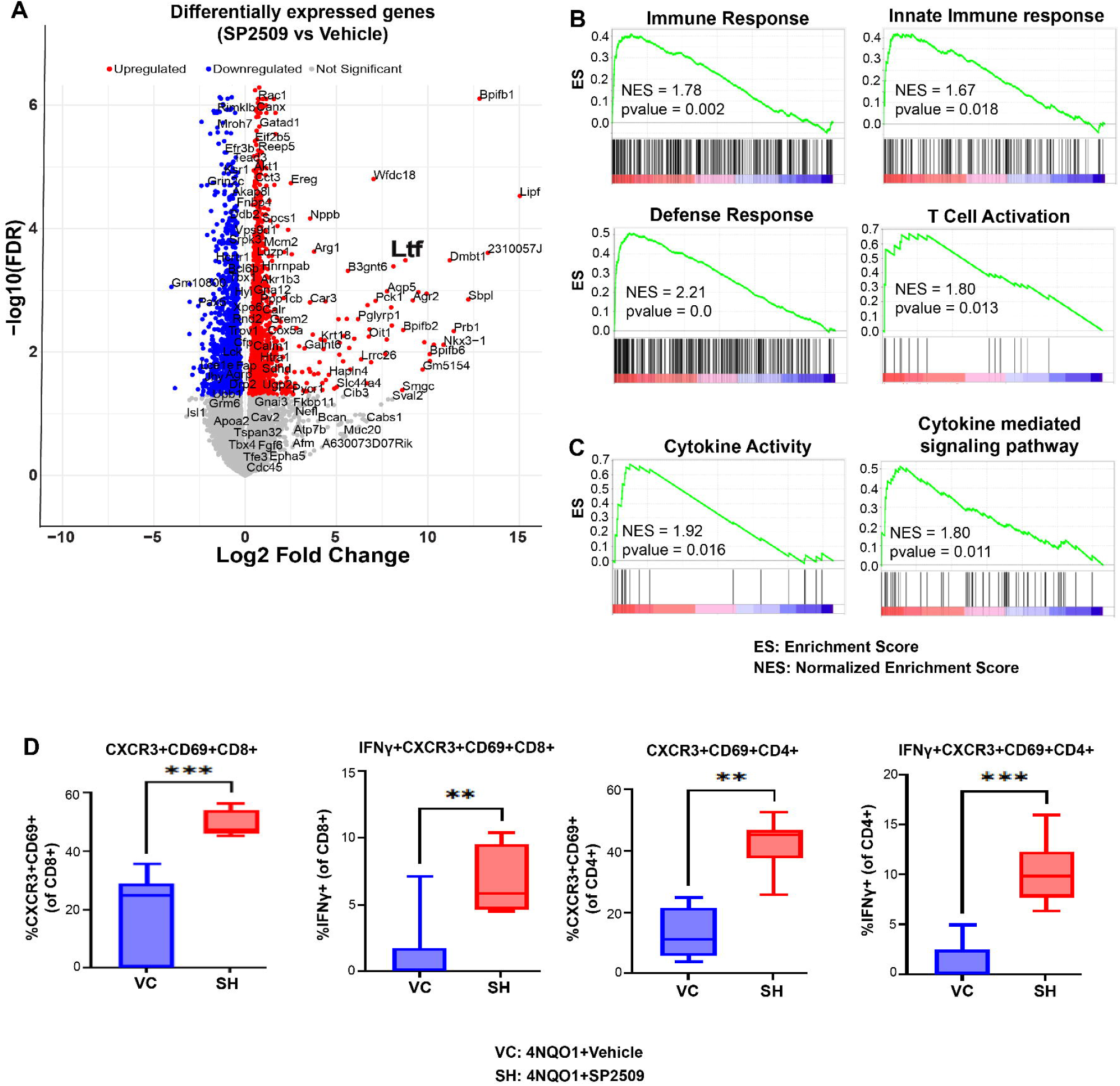
LSD1 inhibition promotes immune response-related gene network in chronic 4NQO-induced progressive HNSCC mouse tongue. A) Volcano plot showing differentially expressed genes between SP2509 vs. Vehicle-treated mice. B) Gene set enrichment analysis (GSEA) of tongue tumors treated with SP2509 showed significant upregulation of innate immune response and activation of T-lymphocytes-related genes as compared to vehicle control. C) GSEA analysis showing increased cytokine activity with SP2509 treatment. D) Flow cytometry data demonstrating CXCR3 and IFNγ in activated CD4+ and CD8+ positive cells. (*p<0.05, **p<0.01, ***p<0.001, ****p<0.0001).

### LSD1 inhibition promotes the IFNγ in T cells, which promotes CXCR3 in T cells and CXCL9 to promote antigen-presenting DCs

We hypothesized that the effects of LSD1 inhibition on CD8+ T cell infiltration and activation are linked to increased antigen presentation activity. To investigate cellular changes in response to LSD1 inhibition, we conducted single-cell RNA-seq (scRNA-seq) experiments. Seurat cluster analysis identified the major cell type compartments (Figure 4A), and focused analysis within the immune compartment revealed that SP2509 treatment promoted the expression of specific DC types (Figure 4B, Figure S4A). The accumulation of DCs was also confirmed by upregulation of the DC-specific marker *Batf3* (Figure 4C, D). Batf3, a critical transcription factor in DCs, is crucial for antigen presentation, cytotoxic T-cell activation, and immunity (40). RT-qPCR analysis also validated the increased overall expression of DC-specific factors Batf3, Cxcl9, and the T cell-related factor Cxcr3 (Figure 4E). Flow cytometric analysis validated that SP2509 promoted the expression of overall DC (Figure 4F) as well as DC subtypes, including conventional DC1 (cDC1), migratory DC (mDC), and resident DC (rDC) (Figure 4G; Figure S4B). CD86 is a well-established activation marker for DCs, which was analyzed in conjunction with a DC-specific marker. SP2509 treatment promoted activated (CD86+) cDC1, mDC, and rDC (Figure 4H). Overall, our data provide strong evidence that SP2509 promotes an increased in activated DC (CD86+) subpopulation. These activated DCs are known to have a role in efficient antigen presentation to T cells (41).

**Figure 4:**
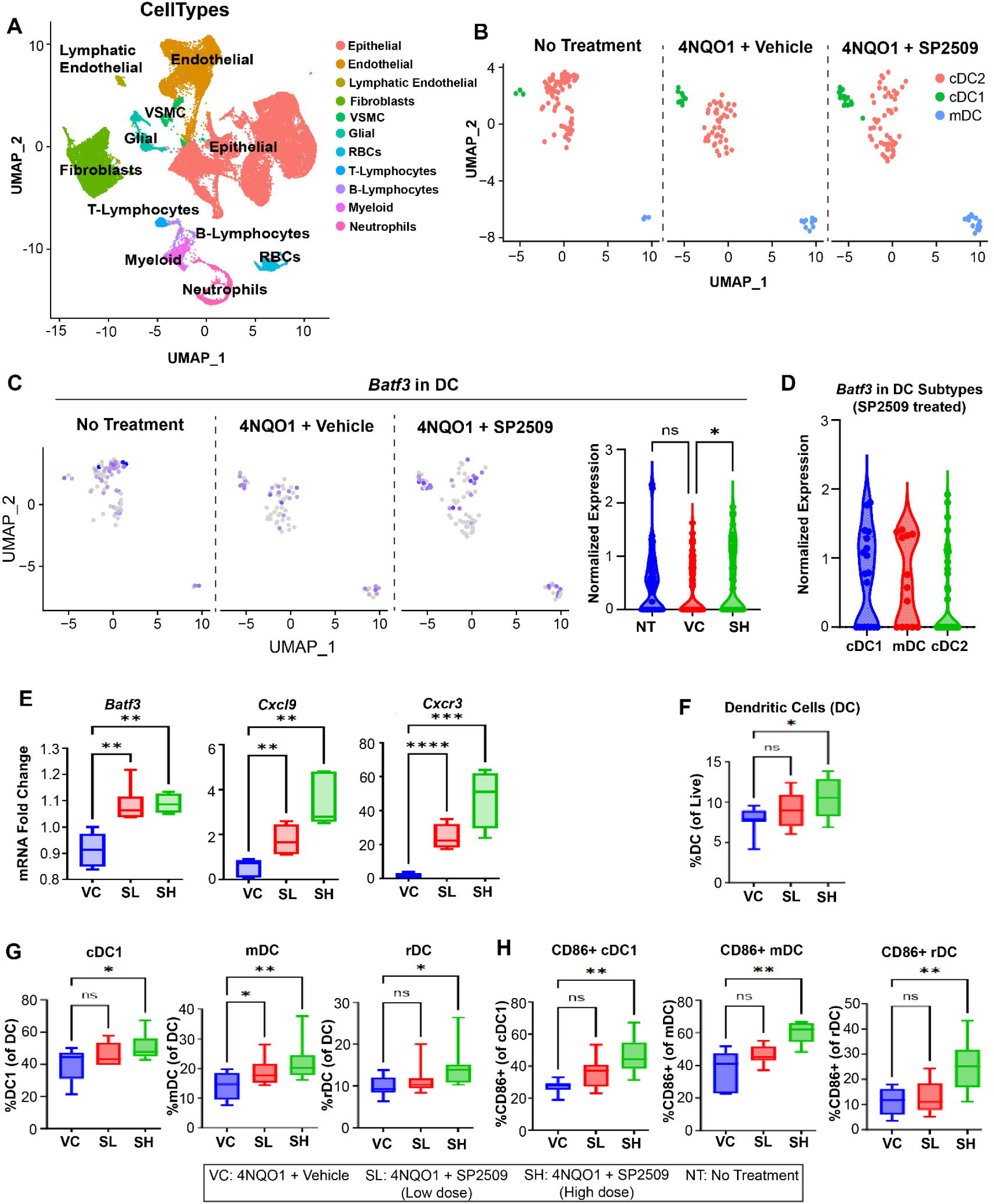
LSD1 inhibition promotes activation and infiltration of DCs in progressive 4NQO HNSCC mice. A) UMAP representation of the cell clusters identified by single-cell RNA sequencing. B) UMAP representation of DC extraction from a pool of total cell clusters. C) Feature plot and violin plot representation of the Batf3 expression in DC with SP2509 treatment. D) Violin plot representation of Batf3 expression in various types of DC with SP2509 treatment. E) RT-qPCR showed a higher expression of Batf3, Cxcr3, and Cxcl9 with SL and SH treatment. F) Box plot representation of the elevated overall DC percentage in the SL and SH treatment groups using flow cytometry. G) Box plot representation of elevated specialized DC percent in SL and SH treatment groups using flow cytometry. H) Box plot representation of activated (CD86+) specialized DC percent in SL and SH treatment groups using flow cytometry. (*p<0.05, **p<0.01, ***p<0.001, ****p<0.0001).

### LSD1 inhibition upregulates MHC class I in the epithelial compartment

Consistently, IPA of the bulk RNA-seq data showed an increase in antigen presentation mediated by MHC-I, dendritic cell migration, and maturation (Figure 5A). To evaluate specific changes in epithelial and immune cells, scRNA-seq data were subjected to chromatin and IPA analysis. Cell chat analysis showed that HNSCC interactions were established in HNSCC, whereas SP2509 promoted interactions and increased the strength of the interactions between DC and T cells (Figure 5B, Figure S4C). Differential epithelial cell expression analysis of HNSCC treated with vehicle vs. SP2509 followed by IPA canonical pathway analysis showed interferon beta and gamma signaling, antigen presentation, and reduced glucose metabolism (Figure S4D). In addition, the differential expression of immune cells in response to SP2509 normalized by vehicle showed Th1 and Th2 pathways, TCR signaling, interferon signaling, and DC maturation (Figure 5C). Interestingly, analysis of specific cell types in scRNA-seq data analysis showed that the SP2509 treatment group had elevated levels of T-cell receptor subunits (TCRβC1 and TCRβC2) (Figure 5D), Cxcl9 in DCs, and Cxcr3 and IFNγ in T cells (Figure 5E; Figure S5A). SP2509 promoted pathways in immune cells and stimulated MHC-I subunits (H2-AA, H2-AB1, H2-Eb1, H2-DMA, and H2-DMB1) (Figure 5F and S5B). Finally, to validate these findings, flow cytometry analysis of MHC-I showed that SP2509 promotes upregulation of MHC-I as compared to vehicle in HNSCC (Figure 5G). Taken together, these results suggest that LSD1 inhibition in HNSCC upregulates MHC class I in epithelial cells, and IFNγ, Cxcl9, and Cxcr3 in immune cells promote T cell and DCs infiltration, representing the likelihood of efficient antigen presentation.

**Figure 5:**
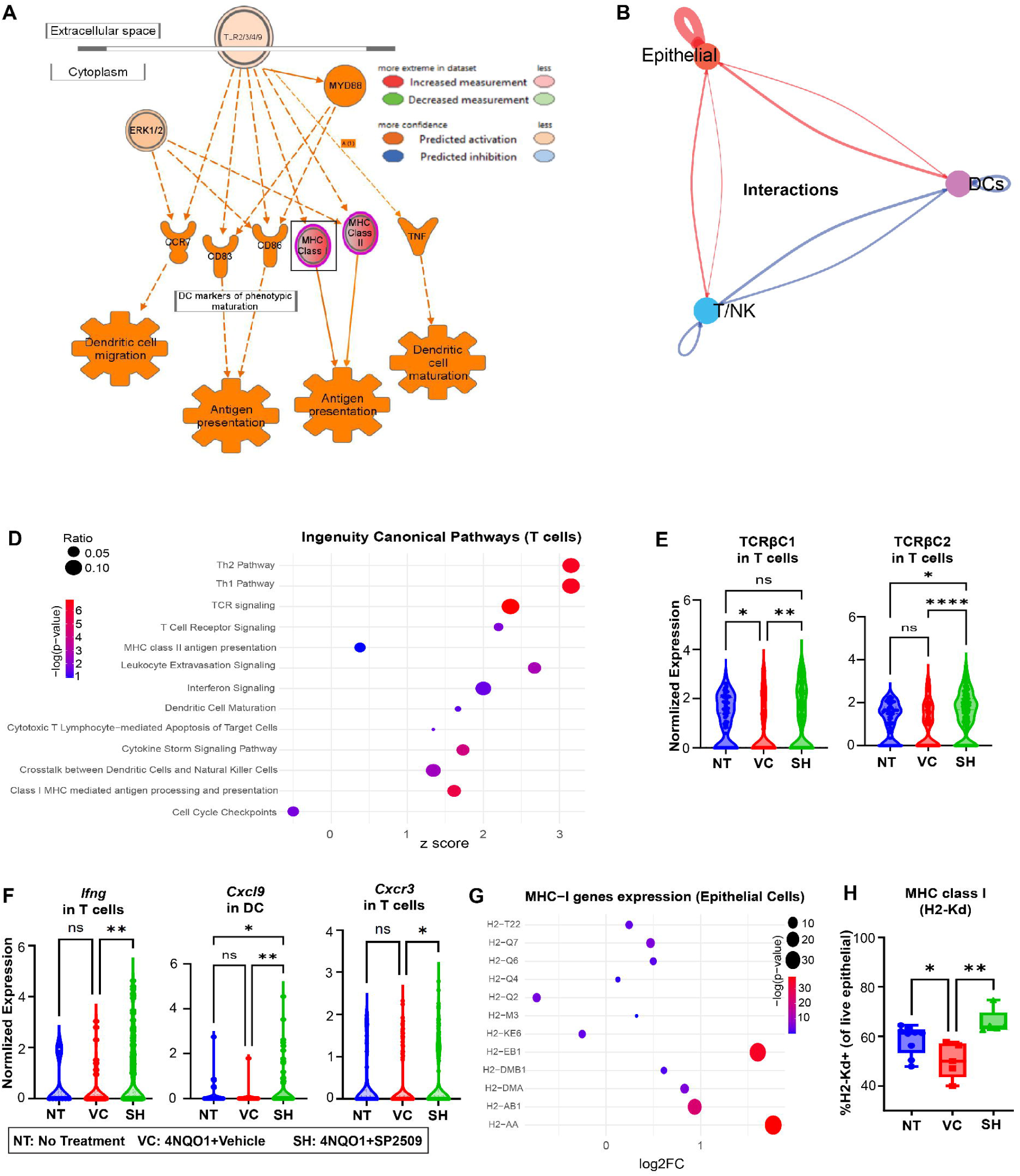
LSD1 inhibition reprograms epithelial cells in HNSCC to upregulate MHC class I and increases cytokine signaling network to promote infiltration of T-cells and DCs. A) IPA of RNA-seq data showing activation of DC migration, antigen presentation, and DC maturation gene network upregulation with SP2509 treatment. B) Cell Chat analysis shows that the directionality of signals originated and its effect on epithelial cells, T cells, or DCs treated in different conditions. A number of interactions were compared when comparing cell-cell interactions in SP2509 treatment vs vehicle in 4NQO mouse tongue via Cell Chat. Blue indicates higher values in SP2509, red indicates higher values in 4NQO. C) Altered pathways within T cells compartment with SP2509 treatment. D) Violin plot showing elevated levels of TCRβ genes in the T cells compartment. E) Violin plots showing increased levels of IFNγ in T cells, cytokines such as Cxcl9 in DCs and Cxcr3 in T cells. F) Altered expression of MHC-I genes with SP2509. G) Percent of MHC class I positive epithelial cells using flow cytometry. scRNA-seq data is presented in violin plot, whereas flow cytometry and RT-qPCR data is presented in box plot. (*p<0.05, **p<0.01, ***p<0.001, ****p<0.0001).

### Humanized HNSCC stem cell mouse model increases CD4, CD8 activation and DC markers after SP2509 treatment

To evaluate the functional effects of LSD1 inhibition in clinical HNSCC specimens, we established a novel humanized HNSCC stem cell model. NOD-Prkdc^em26Cd52^Il2rgem^26Cd22^/NjuCrl (NCG) coisogenic CRISPR genetically engineered immunodeficient mice lack functional T, B, and NK cells, and have mutated SRFα to improve engraftment. NCG mice were humanized by tail vein injection of hPBMC, followed by orthotopic HNSCC stem cell implantation (22,23). These hPBMC-HNSCC stem cells humanized mice treated orally with vehicle or SP2509 for four weeks showed reduced tumor size, coincided with altered tissue shape and pathology, and prevented deterioration of body condition (Figure 6A-C). SP2509 promoted CD4+ and CD8+ T cell infiltration into tongue tumors (p<0.05) (Figure 6D, E), and their activation was determined by the presence of activated CD4+ (CD69+CD4+) and CD8+ (CD69+CD8+) T cells (Figure 6F). RT-qPCR analysis of isolated mRNA showed that SP2509 increased the expression of DC-specific markers Batf3, Cxcr3, and Cxcl9 (Figure 6G). Although there is no perfect model to evaluate the anticancer effects, applying a series of in vivo and humanized HNSCC mouse models validated our findings.

**Figure 6:**
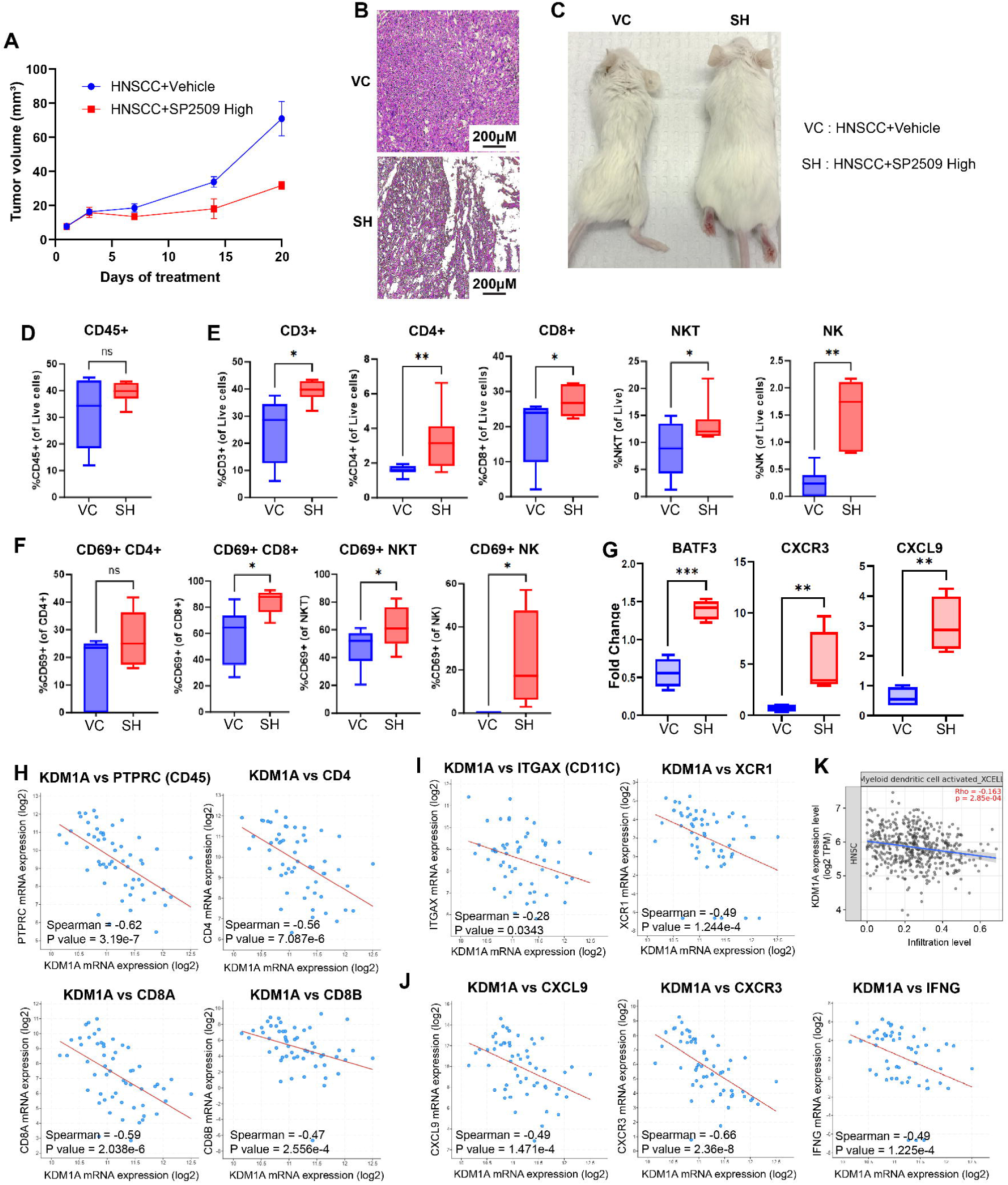
LSD1 inhibition by SP2509 attenuates HNSCC in novel humanized mice model and promotes Batf3, Cxcr3, and Cxcl9 and MHC-I. The cancer genome atlas (TCGA) data showed that Kdm1a has an inverse correlation with immune infiltration-related genes. Humanized mice were generated by first injecting NSG mice with human PBMC (hPBMC), then post 3-days, HNSCC stem cell implantation in the tongue was performed. A) Measurement of tumor volume at 3 to 7-day intervals in vehicle or SP2509 treated mice. B) H&E staining showed that SP2509 attenuated the tumor phenotype. C) Mice treated with SP2509 showed improved overall health compared with the vehicle-treated group. D) Box plot showing nearly unchanged levels of immune cells (CD45+) in SP2509 treated mice tongue compared to vehicle. E) SP2509 treated group showed elevated levels of total T cells (CD3+), including significant infiltration of CD4+ and CD8+ T cells, along with higher levels of NKT and NK immune cells in the mouse tongue. F) Box plots showing activation (CD69+) of CD4+, CD8+, NKT, and NK immune cells in SP2509 treated mice tongue. G) RT-qPCR showing higher expression of Batf3, Cxcr3, and Cxcl9 with SP2509 high and low dose treatment. The cancer genome atlas (TCGA) data showed that KDM1A has an inverse correlation with immune infiltration-related genes, including H) immune cell marker gene CD45 (PTPRC), CD4, and CD8B T cells, I) DC cell marker genes ITGAX (CD11c) and XCR1, and J) Cytokine genes CXCL9, CXCR3, and IFNγ. K) Negative correlation of DCs infiltration with KDM1A expression. (*p<0.05, **p<0.01, ***p<0.001, ****p<0.0001).

### KDM1A has an inverse correlation with IFNγ, CXCR3, and CXCL9 in clinical tumors

To evaluate whether the findings from murine and humanized HNSCC models were relevant to human tumor samples, human HNSCC patient data from The Cancer Genome Atlas (TCGA) were analyzed. We categorized patients as human papillomavirus (HPV)-negative (Figure 6H-J, Figure S6A-C) and HPV-positive (Figure S6D-F) and subjected them to individual groups for the correlation analysis. In HPV-negative HNSCC patients, KDM1A was significantly inversely correlated with the hematopoietic cell marker gene PTPRC (CD45) and the T cell marker genes CD4, CD8A, and CD8B (Figure 6H). We also found that KDM1A mRNA expression was negatively correlated with the specific DC markers ITGAX (CD11C) and XCR1 (Figure 6I). The DC-specific chemokine CXCL9, CD8-specific chemokine receptor CXCR3, and CD8 cell-activating cytokine IFNG were also inversely correlated with KDM1A expression (Figure 6J). Additionally, immune infiltration analysis revealed a significant negative correlation between KDM1A expression and activated myeloid DCs (Figure 6K).

### HNSCC epithelial cells are reprogrammed by LSD1 knockout, or inhibition secretes IFNγ to stimulate MHC-I in HNSCC as well as CXCR3 in CD8 and CXCL9 in DCs

To evaluate whether HNSCC cells were reprogrammed to upregulate MHC-I and signal immune cells, we analyzed Krt14 promoter-specific *Kdm1a* deletion (*Kdm1a*^-/-^) mouse tongues and co-cultured immune cells after knockdown of Kdm1a in HNSCC cells. Our earlier studies showed that *Kdm1a* knockout in epithelial cells attenuates HNSCC and inhibits the expression of key oncogenes (2). Immunofluorescence analysis of *Kdm1a*^-/-^ HNSCC tongues showed upregulation of TCRβ, MHC-I, and CD8+ T cells compared to tongues from *Kdm1a*^WT/WT^ mice (Figure 7A, Figure S7A). In addition, SP2509 treatment resulted in the upregulation of TCRβ, MHC-I, and CD8+ T cells compared to the tongue of vehicle-treated mice (Figure 7B, Figure S7B).

**Figure 7:**
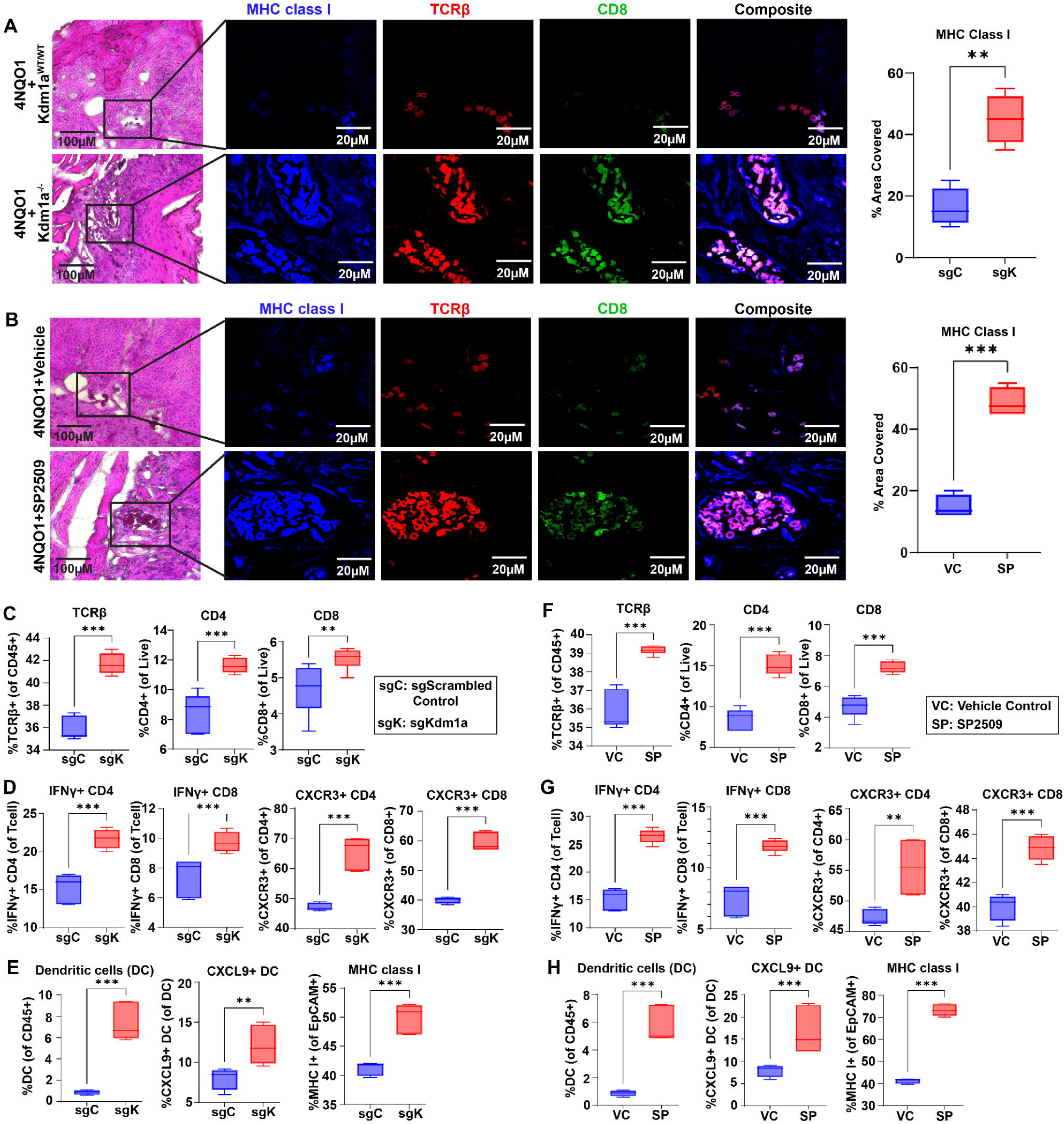
Deletion of Kdm1a in Krt14-expressing HNSCC mice tongue, KDM1A sgRNA treatment or SP2509 treatment in co-culture increases MHC-I+, TCRβ+ and CD8+ cells. A) Tongue epithelial cells specific deletion of Kdm1a shows increased expression of MHC-I, TCRβ and CD8+ cells compared to 4NQO1-induced mouse HNSCC tongue tumors. B) SP2509 treatment increased expression of MHC-I, TCRβ+, and CD8+ immune cells in 4NQO1-induced mouse HNSCC tongue tumors. C) KDM1A sgRNA treatment followed by flow cytometry analysis of KDM1A sgRNA treated co-culture model shows an increase in TCRβ+, CD4+, and CD8+ cells. D) IFNγ+CD4+, IFNγ+CD8+, CXCR3+CD4+ and CXCR3+CD8+ T cell population. E) DCs (CD11c+ MHC class II+), CXCL9+ DCs and MHC-I+ Epcam+ cancer epithelial cells. F) SP2509 treatment followed by flow cytometry analysis in HSC3:hPBMC co-culture model shows an increase in TCRβ+, CD4+, and CD8+ cells G) IFNγ+CD4+, IFNγ+CD8+, CXCR3+CD4+ and CXCR3+CD8+ T cell population H) DCs (CD11c+ MHC class II+), CXCL9+ DCs and MHC-I+ Epcam+ cancer epithelial cells. (*p<0.05, **p<0.01, ***p<0.001, ****p<0.0001).

To evaluate whether HNSCC cancer cells are reprogrammed to secrete factors that stimulate CD8+ T cells and DCs, we established a co-culture model of HSC3 (OSCC cell line) and human peripheral blood mononuclear cells (PBMCs). First, HSC3 cells were transfected with *Kdm1a* sgRNA construct or scrambled control for 24 h, washed several times, and incubated with PBMCs for another 48 h to observe the effect on specific immune cells, chemokines, and cytokines. Both *Kdm1a* knockout and LSD1 inhibition resulted in a significant difference in TCRβ, MHC-I, and CD8+ T cells compared to their respective controls (Figure S7C-D). Flow cytometry analysis revealed increased TCRβ+, CD4+, and CD8+ T cells and an increased DC population (CD11c+ MHC class I+) (Figure 7C, E) in the *Kdm1a* knockout group. HNSCC cell co-culture with human PBMCs showed that *Kdm1a* knockout promoted IFNγ production in CD4+ and CD8+ T cells, DCs, CXCL9 in DCs, and MHC-I in CD326+ cells (Figure 7D, E).

Next, the HSC3 cells (OSCC cell line) were treated with SP2509 for 24 h, washed with PBS to remove residual SP2509, and incubated with PBMCs. The results showed that SP2509 caused a statistically significant increase in TCRβ+, CD4+, and CD8+ T cells (Figure 7F), IFNγ production in CD8+ and CD4+ T cells, CXCR3 production in CD4+ and CD8+ T cells, as well as in DC, CXCL9+ DC, and MHC-I in CD326+ cells (Figure 7G-H). Overall, these results show that LSD1 inhibition and *Kdm1a* genetic deletion reprogram HSC3 cells to upregulate MHC class I and secrete IFNγ, which stimulates the CXCL9-CXCR3 axis and antigen-presenting DCs, and increases cytotoxic T cell production, leading to antitumor immunity.

To evaluate changes in immune cells and their activation through the CXCL9-CXCR3 pathway, we performed blocking experiments with anti-human CXCL9 and anti-human CXCR3 as shown in previous studies (42,43). HSC3 and human PBMCs co-culture studies showed significantly (p<0.001) reduced proliferation of immune cells (CD45+), CD4+ and CD8+ T cells, DCs, and production of IFNy after treatment with anti-human CXCL9 and anti-human CXCR3 blocking antibodies (Figure 8A-D).

**Figure 8:**
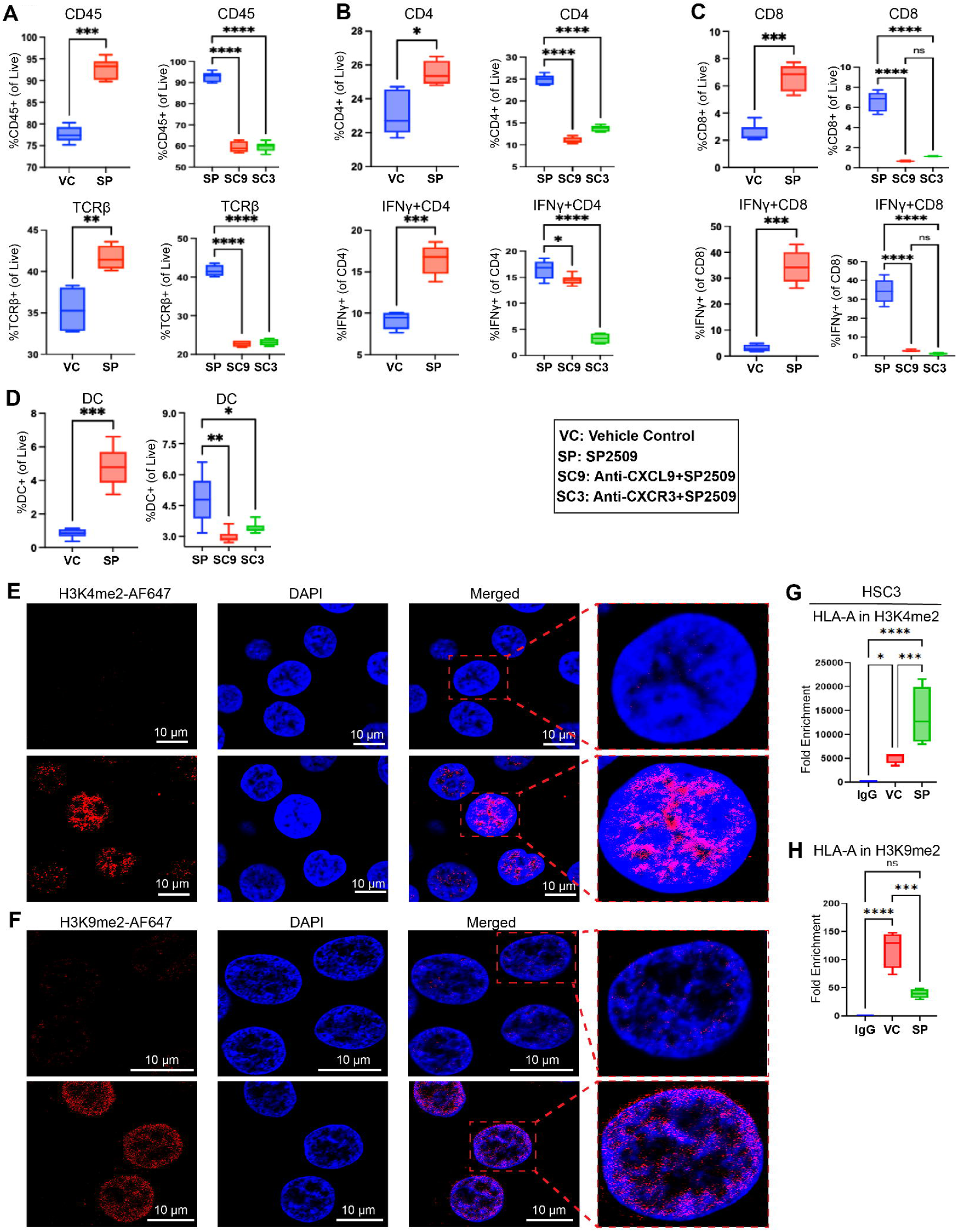
LSD1 regulates IFNγ production through CXCL9-CXCR3 axis and regulates MHC class I gene expression through histone H3K4 and H3K9 modifications. A) Major immune cell marker CD45 and T cell marker TCRβ got reduced significantly after blocking with CXCL9 and CXCR3 antibodies in respective to LSD1 inhibited group. B-C) CD4+ T cells, CD8+ T cells and their activation and cytotoxic effect through IFNγ production were significantly reduced after CXCL9 and CXCR3 blocking in respect to LSD1 inhibited group. D) DC (CD11c+ MHC class II+) population was also reduced after CXCL9 and CXCR3 blocking. E-F) Amount of H3K4me2 and H3K9me2 were visibly increased after LSD1 inhibition. G-H) MHC class I gene HLA-A enrichment was significantly increased in H3K4me2, whereas significantly decreased in H3K9me2. (*p<0.05, **p<0.01, ***p<0.001, ****p<0.0001).

### LSD1 regulates H3K4me2 and H3K9me2 on MHC class I gene

To evaluate the changes in H3K4 and H3K9 methylation status by LSD1, immunofluorescence analysis of HSC3 was performed. The ananlysis showed increased H3K4me2 and H3K9me2 levels with LSD1 inhibition (Figure 8E). A recent study showed that LSD1 acts via an autoregulatory mechanism (44). In order to evaluate changes in chromatin, chromatin immunoprecipitation (ChIP) with H3K4me2 and H3K9me2, we evaluated the status of these di-methylated histones on the MHC class I gene HLA-A and Kdm1A. The data showed that n significantly (p<0.0001) upregulated enrichment of HLA-A in H3K4me2 and downregulated enrichment in H3K9me2 (p<0.001) by LSD1 inhibitor as compared to the vehicle control (Figure 8F). We also have observed significant (p<0.0001) changes of KDM1A in H3K4me2 while KDM1A in H3K9me2 was not changed significantly (Figure S8E-F) suggesting a probable self-regulatory function of LSD1.

## Discussion

Increasing evidence indicates that HNSCC evolution involves dynamic interactions among different cell subpopulations that are driven, in part, by epigenetic changes in tumor cells and their surrounding microenvironment. This study aimed to investigate the role of LSD1 inhibition in promoting the activation of CD8+ T cells and inducing antitumor immunity. Our previous studies have shown that aberrant LSD1 activation leads to HNSCC progression (1,2). LSD1 irreversible inhibitor Bomedemstat has been shown to inhibit small-cell lung carcinoma (SCLC) (45). LSD1 inhibition has also been shown to upregulate MHC-I expression in SCLC (46). LSD1 regulates gene expression, DNA demethylation, and CD8+ T cell upregulation (47). The gap in understanding the mechanism of infiltration and activation of CD8+ T cells and MHC-I upregulation has not been investigated. We identified novel mechanisms of LSD1 in 1) DC-mediated antigen presentation via the IFNγ-CXCL9-CXCR3 axis and 2) reprogramming of HNSCC cells to promote and upregulate MHC-I molecules in vivo in murine and humanized models relevant to clinical HNSCC.

MHC class I peptides, such as HLA-A or HLA-B in humans and H2-Q7 or H2-Eb1 in mice, interact with T-cell receptor β (48–50). High levels of sustained IFNγ production by TH1 cells or CTLs typically require T-cell receptor (TCR)-mediated recognition of antigens in the context of MHC class I molecules (51). C-X-C motif chemokine 9 (CXCL9), CXCL10, and CXCL11 are produced in response to interferon-γ (IFNγ) and trigger inflammation by accumulating activated lymphocytes (52,53). LSD1 inhibition also upregulates a coordinated network of IFNγ, CXCL9, and CXCR3, which are required for T-cell infiltration and antitumor immunity. In the tumor microenvironment, these chemokines help establish a “hot” tumor microenvironment by initiating a chemokine network (54).

Furthermore, our study showed that LSD1 inhibition promoted IFN-γ-expressing T cells. IFNγ-inducible CXCL9 produced by cells recruits CXCR3-expressing CD8+ T cells from the circulation into the tumor bed (55,56) and could lead to a T cell-inflamed (hot) tumor microenvironment (42). A recent study has shown that CD4+, CD8+, and DCs act as a triad to promote antitumor immunity(4), which is consistent with our finding that LSD1 inhibition promotes CXCL9 expression in DCs, CD4+, and CD8+ cells. Newly recruited CXCR3+ T cells produce IFNγ in response to antigen engagement, thereby inducing the additional production of CXCL9 by DCs, leading to increased T cell infiltration. Thus, we provide functional evidence that LSD1 inhibition enhances DC-mediated T-cell activation and increases the DC-specific marker Batf3. Batf3, a transcription factor essential for the development of cDC1s, is linked to the production of CXCL9 and CXCL10 in the tumor microenvironment (TME) (57). Overall, understanding the role of LSD1 could be a new player in promoting antigen-presenting DCs via the coordinated network IFNγ-CXCL9-CXCR3 to activate CD8+ T cells to induce antitumor immunity.

We also found that Kdm1a expression was significantly inversely correlated with T cell-related genes (CD4, CD8, and CD3), DC-related genes (CXCL9 and XCR1), CD8+ cell activation markers (CXCR3), and T cell-induced factors (IFN-γ), whereas Kdm1a inhibition increased their expression in functional studies on syngeneic, chronic tobacco carcinogens, and humanized HNSCC stem cell models.

Although this is the first study to evaluate the role of LSD1 in DC-mediated antitumor immunity, few studies have investigated its underlying mechanisms. The stimulation of tumor suppressors or attenuation of oncogenic proteins can induce antitumor immunity, which includes activation of the P53 (58) melanocortin-1 receptor (MC1R) (59) EZH2 mediated suppression (60) trans-vaccenic acid diet (61), and LSD1 inhibition to promote endogenous retrovirus expression (62).

We showed that LSD1 inhibition reprogrammed HNSCC cells to upregulate mouse MHC-I molecules, as well as human MHC-I, as in humanized mouse models. Loss or downregulation of MHC-I in cancer cells is a major mechanism underlying resistance to T cell-based immunotherapies. These have been shown in studies of TRAF3 inhibition (63), chemoresistance, and immunotherapy resistance mechanisms (63). Histone methyltransferase WHSC1 (64). H3K4me2 is present in the vicinity of active genes (PMID: 15680324). Our study provides new insights into the role of LSD1 in promoting MHC-I by promoting H3K4me2 activation and direct interaction with MHC-I. Overall, these findings could have important implications for the development of new cancer immunotherapies.

In conclusion, we demonstrated for the first time that LSD1 inhibition promotes antigen-presenting DCs to activate CD8+ T cells to induce antitumor immunity. We also found that LSD1 inhibition promoted the IFNγ-CXCL9-CXCR3 axis for the activation of DCs and MHC-I upregulation in HNSCC tumor cells. Finally, we identified a novel mechanism by which LSD1 inhibition promotes the activation of H3K4me2 and its direct interaction with MHC-I. These LSD1-mediated mechanisms have an impact on epigenetic therapy, resistant HNSCC therapies, and poorly immunogenic tumors.

## Acknowledgments

Support: This study was supported by NIH grants R01 DE031413 (MB), R01 DE030350 (MAK, SM, and XV), R01 DE033519 (XV, SM, and MAK), R01 DE030350 S1 (SM), R01 DE 031831 (SM) and CTSA pilot grant UL1TR001430 (MB). We are also thankful to the Flow Cytometry Core Facility of Boston University and the Microarray Core Facility of Boston University.

## Author contributions

Amit Kumar Chakraborty, Lina Kroehling, Rajnikant Dilip Raut, Chumki Choudhury, and Manish V. Bais performed the experiments and analyzed the data. Amit Kumar Chakraborty, Bikash Sahay, Maria Kukuruzinska, Silvio Gutkind, Xaralabos Varelas, Stefano Monti, and Manish V. Bais helped with the conception, interpretation, and manuscript editing. Manish Bais conceived of and designed the study, interpreted the data, and wrote the manuscript.

---

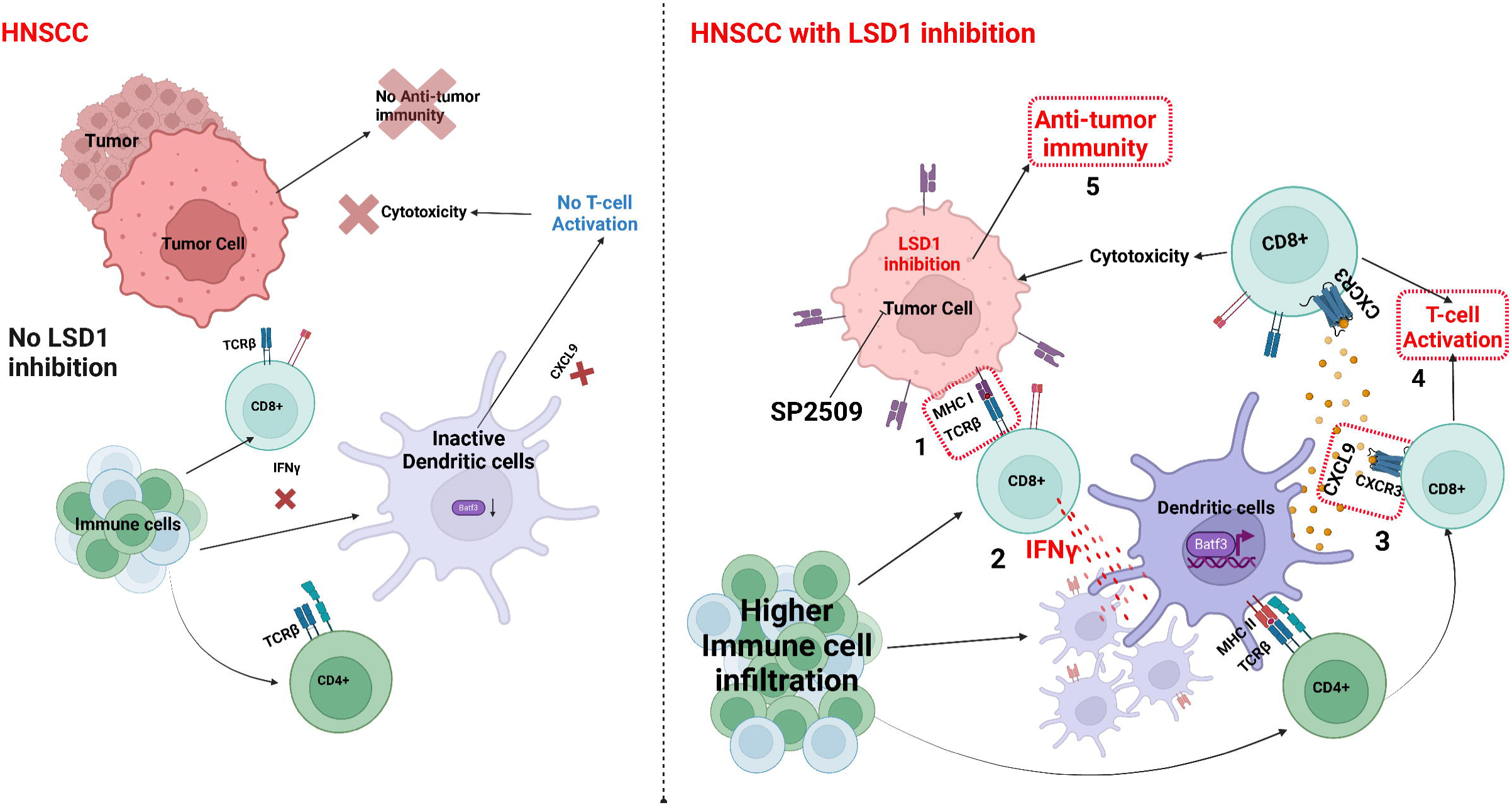

