## Supplementary material for "LSD1 inhibition corrects dysregulated MHC-I and dendritic cells activation through IFNγ-CXCL9-CXCR3 axis to promote antitumor immunity in HNSCC": Figure S1

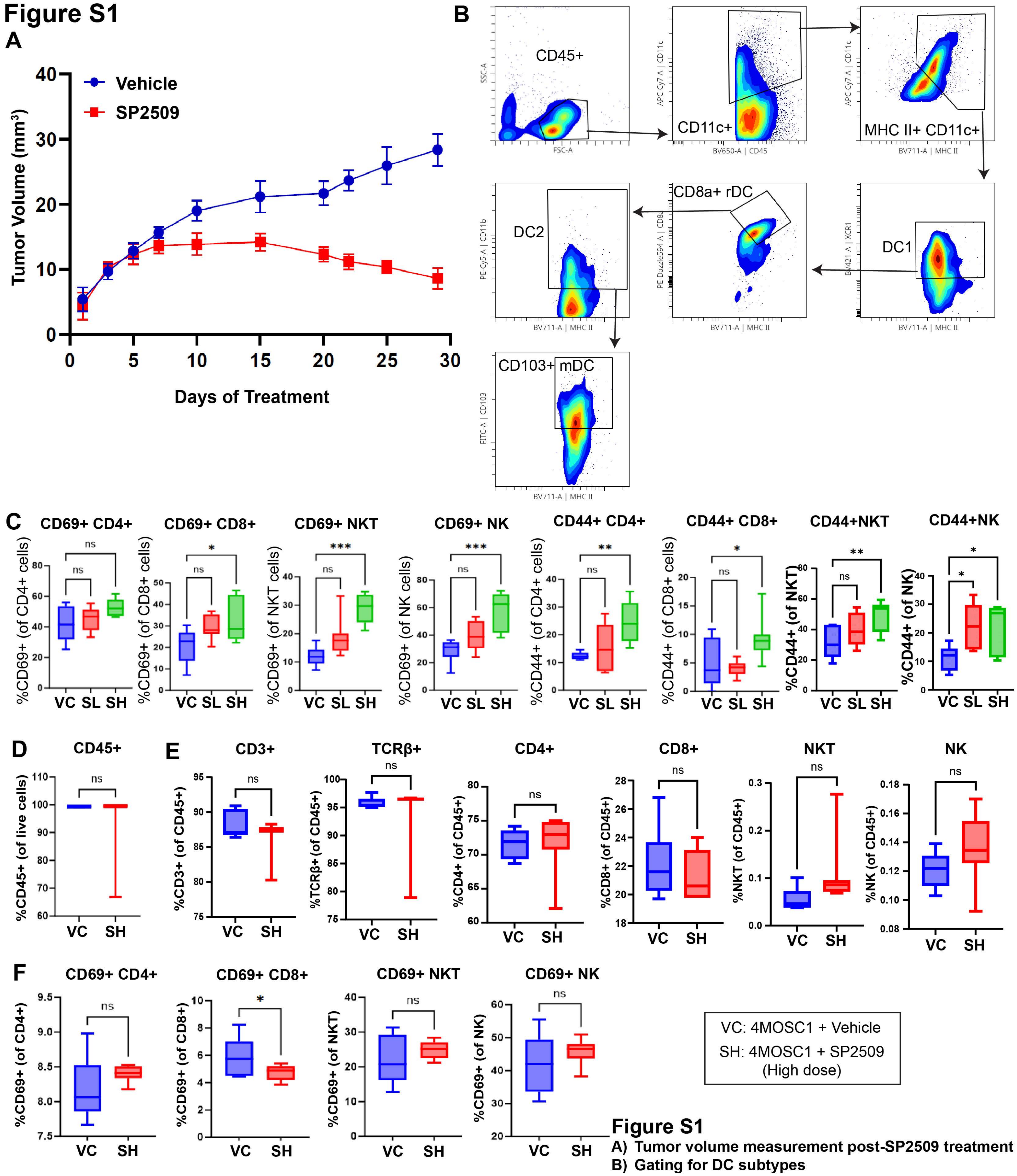


**Figure S1**

A) Tumor volume measurement post-SP2509 treatment
B) Gating for DC subtypes
C) Activated T cells
D) Total immune cell popupation (CD45+) in spleen
E) Quantification of T cells and NK cells in spleen
F) Activation status of T cells and NK cells in spleen
