## Supplementary material for "LSD1 inhibition corrects dysregulated MHC-I and dendritic cells activation through IFNγ-CXCL9-CXCR3 axis to promote antitumor immunity in HNSCC": Figure S2

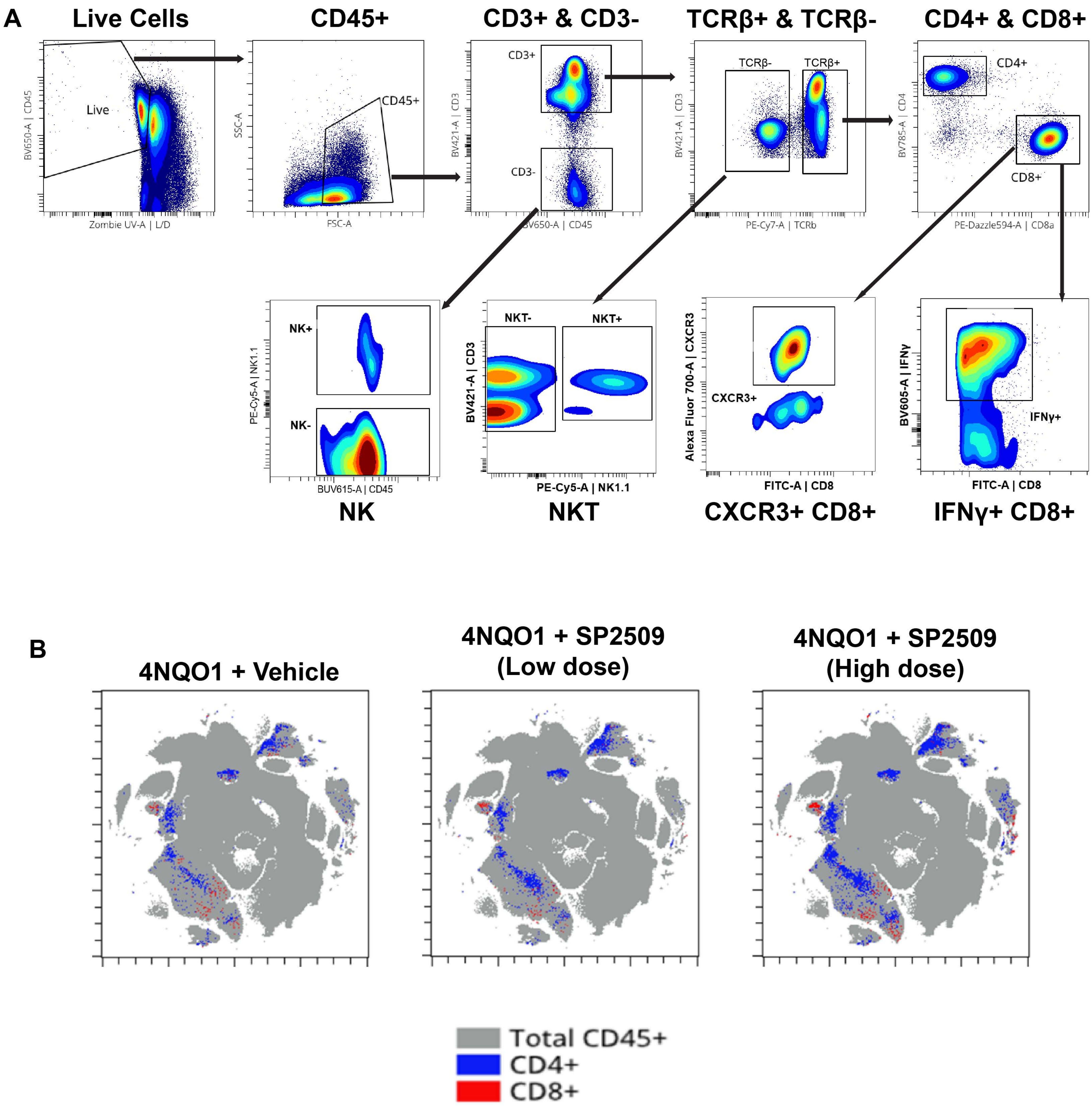


**Figure S2**

A) Gating strategy for the analysis of immune cell population in Vehicle and SP2509 treated 4NQO1 mice model.

B) t-SNE plots showing population of CD4+ and CD8+ T cells.
