## Supplementary material for "LSD1 inhibition corrects dysregulated MHC-I and dendritic cells activation through IFNγ-CXCL9-CXCR3 axis to promote antitumor immunity in HNSCC": Figure S3

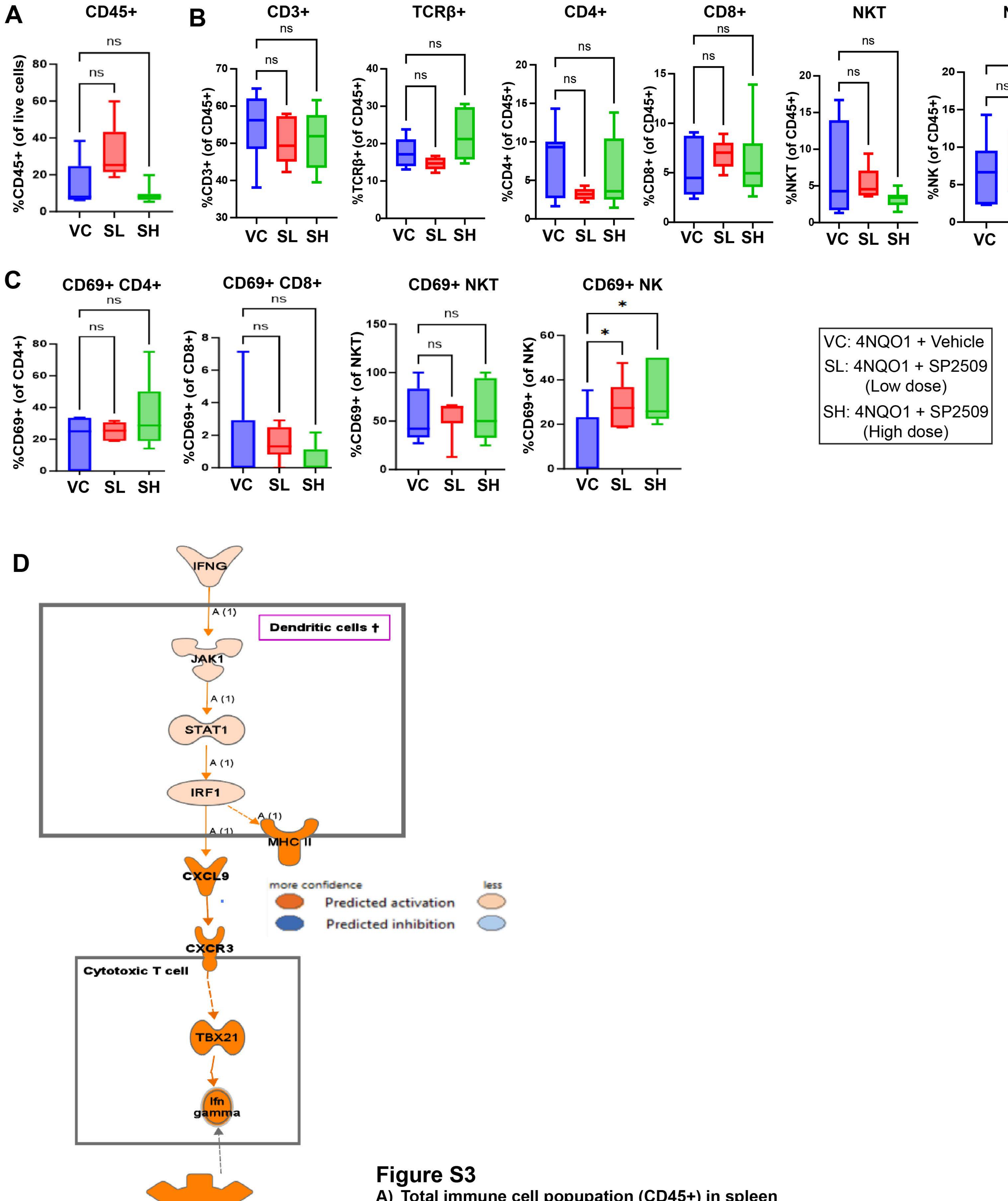


**Figure S3**

A) Total immune cell popupation (CD45+) in spleen
B) Quantification of T cells and NK cells in spleen
C) Activation status of T cells and NK cells in spleen
D) RNA-seq followed by IPA shows the IFNγ activation via CXCL9 and CXCR3 axis.
