## Supplementary material for "LSD1 inhibition corrects dysregulated MHC-I and dendritic cells activation through IFNγ-CXCL9-CXCR3 axis to promote antitumor immunity in HNSCC": Figure S4

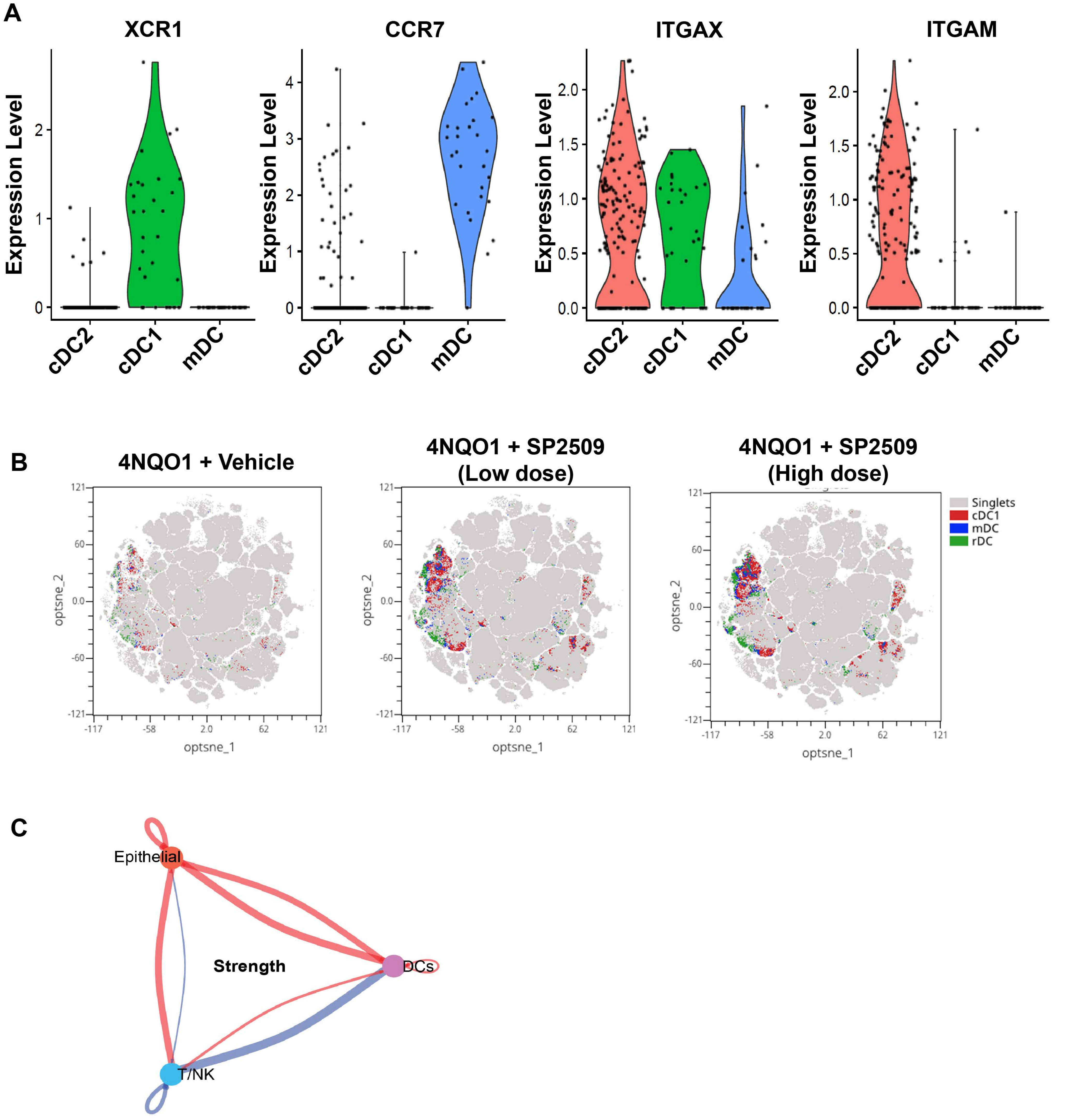


**Figure S4**

**A) Cell Markers for cDC1, cDC2, and mDC in scRNA-seq analysis.**
**B) t-SNE plots showing population of subtypes of dendritic cells using flow cytometry.**
**C) Interaction strength between DCs, epithelial cells, and T/NK cells from CellChat analysis.**
