## Supplementary material for "LSD1 inhibition corrects dysregulated MHC-I and dendritic cells activation through IFNγ-CXCL9-CXCR3 axis to promote antitumor immunity in HNSCC": Figure S5

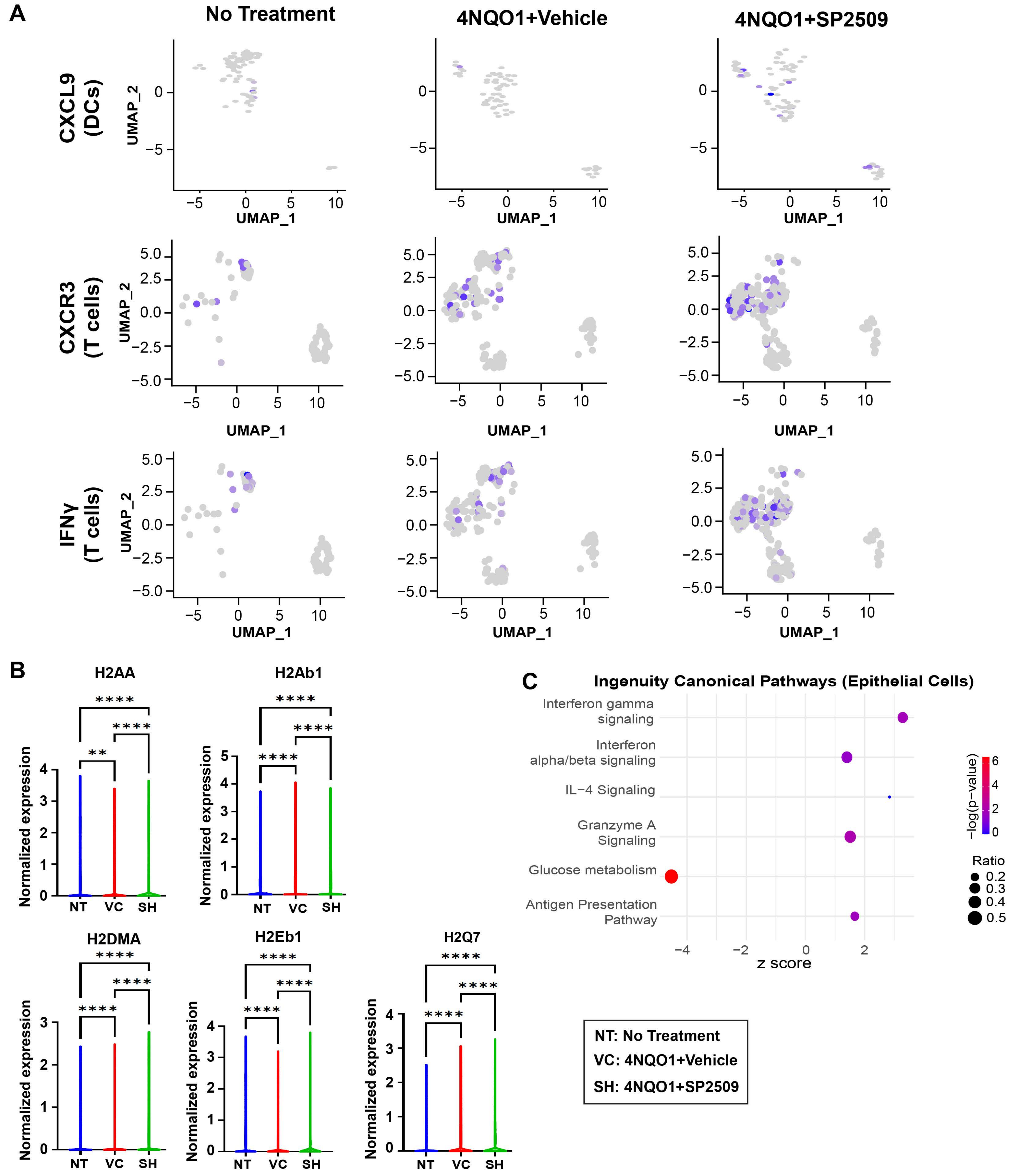


**Figure S5**

A) UMAP plots showing population of cells with the expresion of cytokines.
B) Violin plots showing expression of MHC class-I genes.
C) Altered pathways within epithelial cells compartment with SP2509 treatment
