## Supplementary material for "LSD1 inhibition corrects dysregulated MHC-I and dendritic cells activation through IFNγ-CXCL9-CXCR3 axis to promote antitumor immunity in HNSCC": Figure S6

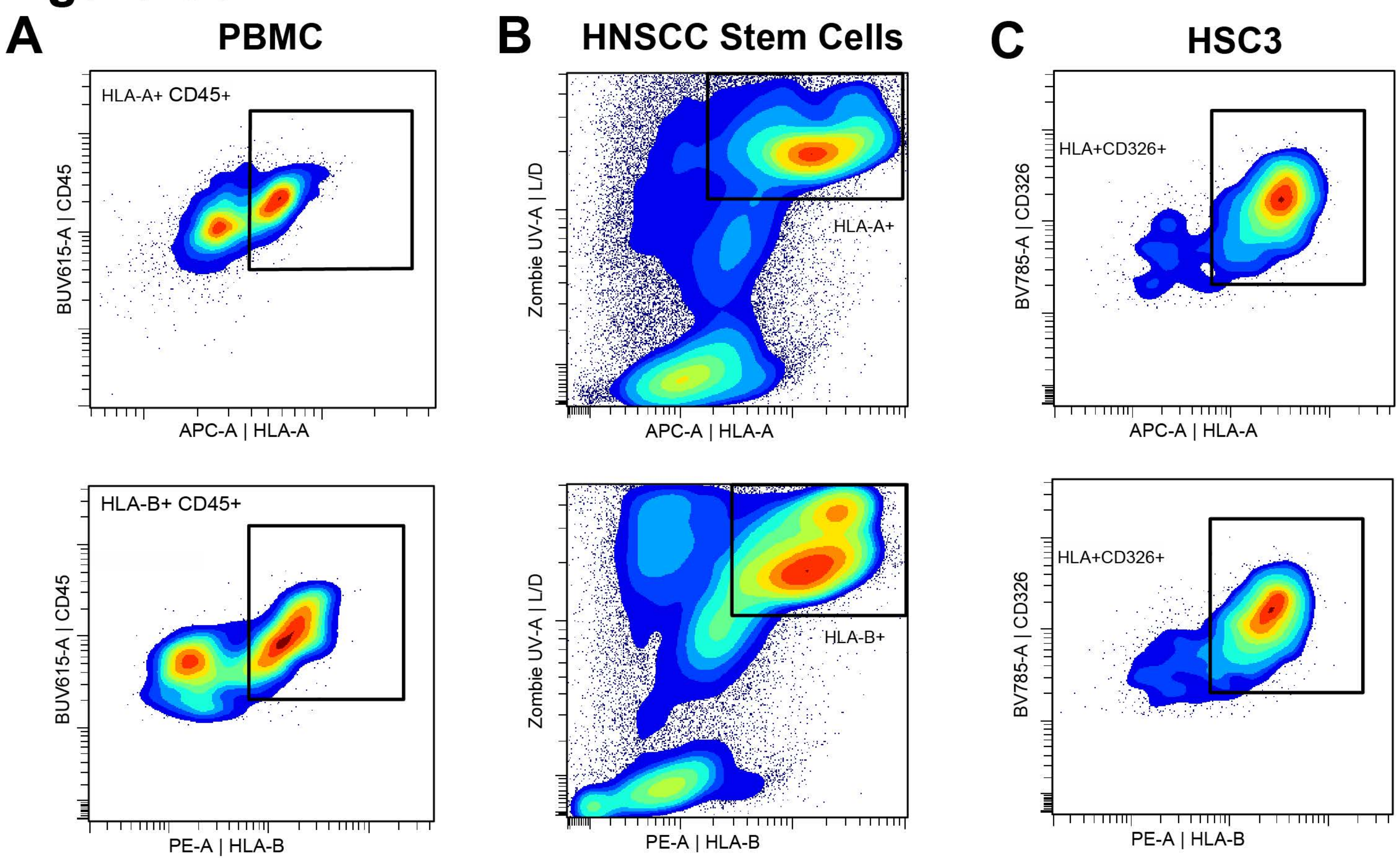


E

**HNSCC Stem Cell Markers**

| Marker Name | Allele Size |
|---|---|
| Amelogenin | X |
| CSF1PO | 12 |
| D13S317 | 8, 11 |
| D16S539 | 12, 13 |
| D5S818 | 13 |
| D7S820 | 10, 12 |
| TH01 | 7, 9.3 |
| TPOX | 8, 11 |
| vWA | 18, 19 |

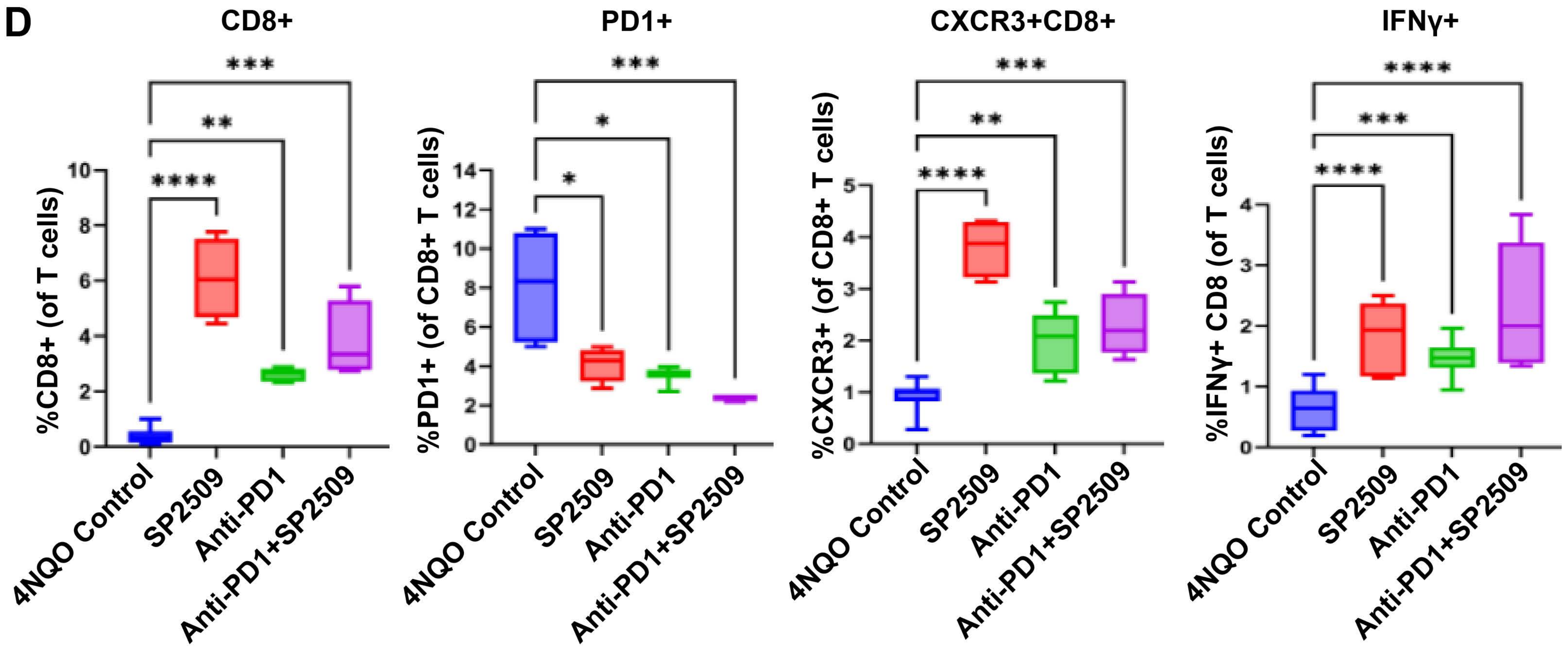


**Figure S6**

**A-C) Results of HLA matching in human PBMC, HNSCC stem cells, and HSC3 cells. Both HLA-A and HLA-B are present.**
**D) Cytotoxic CD8+ T cell profile after Anti-PD1 and combination therapy.**
**E) Table for HNSCC stem cell positive markers.**
