## Supplementary material for "LSD1 inhibition corrects dysregulated MHC-I and dendritic cells activation through IFNγ-CXCL9-CXCR3 axis to promote antitumor immunity in HNSCC": Figure S7

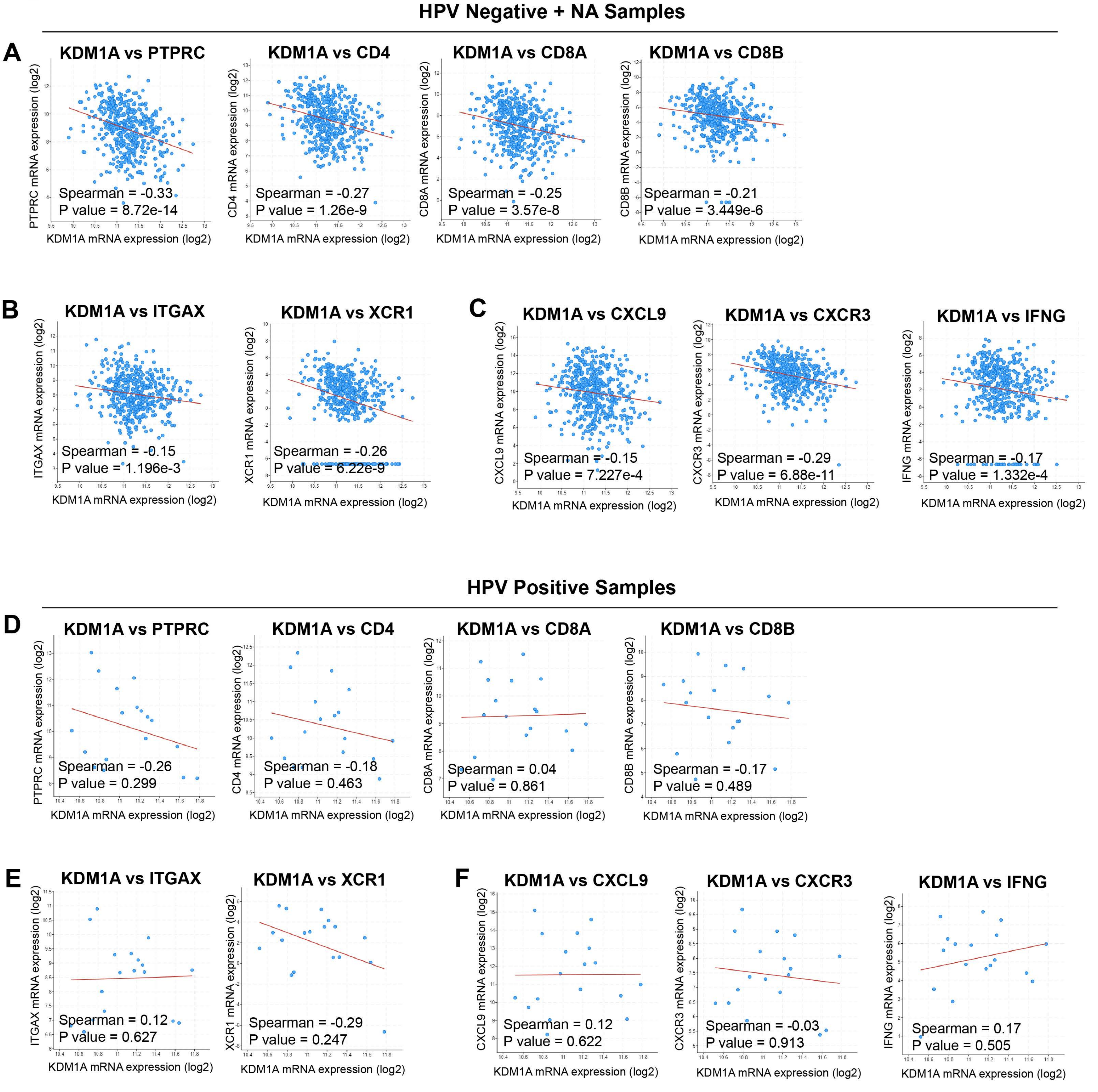


**Figure S7**

**A-C) Spearman correlation of immune genes with KDM1A in HPV- and patients with undisclosed HPV status.**

**D-F) Spearman correlation of immune genes with KDM1A in HPV+ patients.**
