## Supplementary material for "LSD1 inhibition corrects dysregulated MHC-I and dendritic cells activation through IFNγ-CXCL9-CXCR3 axis to promote antitumor immunity in HNSCC": Figure S8

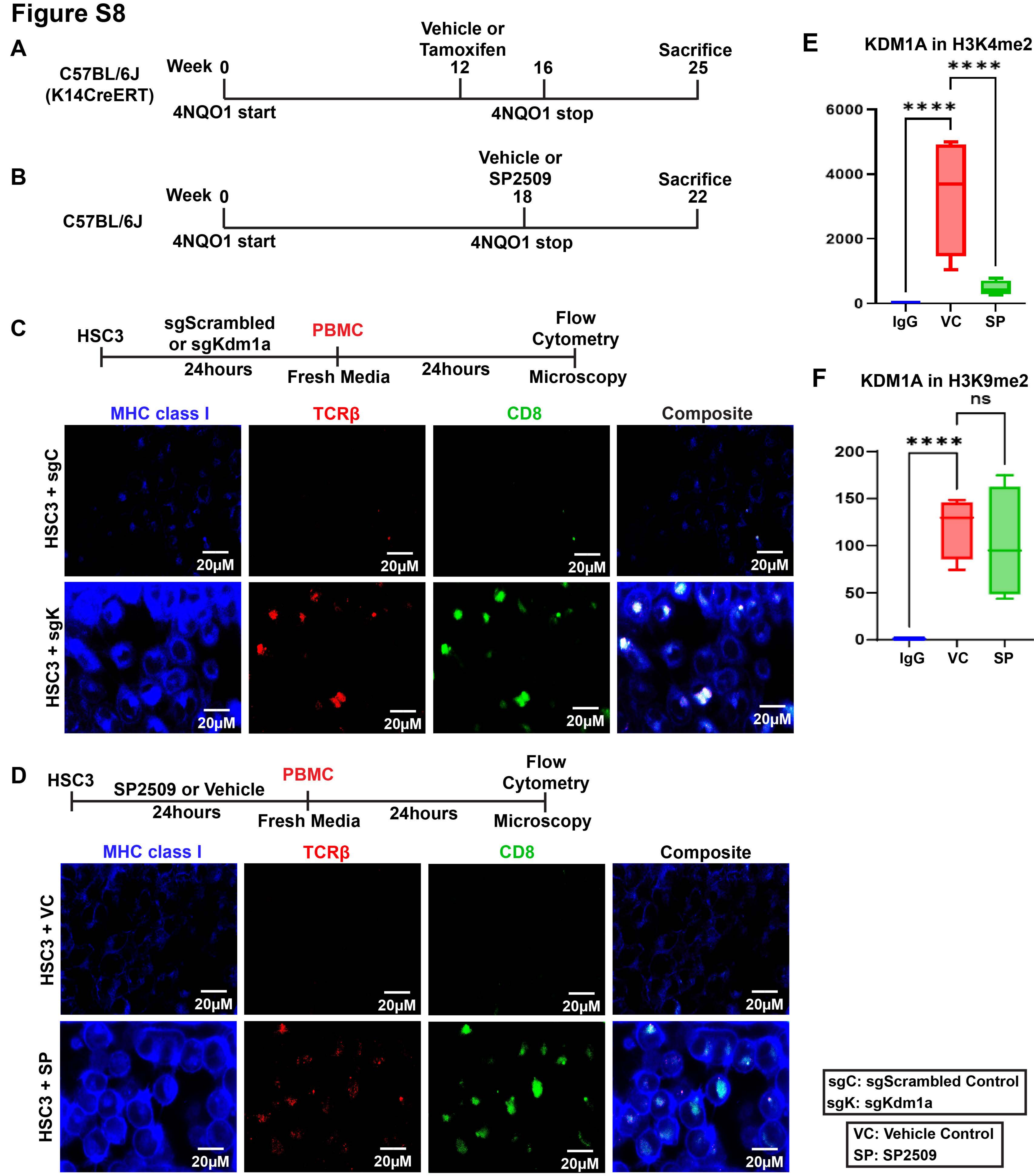


**Figure S8**

A) Experimental strategy for Kdm1a knockout mice.

B) Experimental strategy for phamacological inhibition of LSD1 (Kdm1a).

C) Experimental strategy and visualization of MHC-I, TCRβ and CD8 expressiomn using confocal microscopy in Kdm1a knockout HSC3 cells.

D) Experimental strategy and visualization of MHC-I, TCRβ and CD8 expressiomn using confocal microscopy in SP2509 treated HSC3 cells.

E-F) Fold change of KDM1A in H3K4me2 and H3K9me2.
